# Pathway dynamics can delineate the sources of transcriptional noise in gene expression

**DOI:** 10.1101/2020.09.30.319814

**Authors:** Lucy Ham, Marcel Jackson, Michael P.H. Stumpf

**Affiliations:** School of BioSciences, University of Melbourne, Australia; Department of Mathematics and Statistics, La Trobe University, Australia; School of Mathematics and Statistics, University of Melbourne, Australia

## Abstract

Single-cell expression profiling has opened up new vistas on cellular processes. Among other important results, one stand-out observation has been the confirmation of extensive cell-to-cell variability at the transcriptomic and proteomic level. Because most experimental analyses are destructive we only have access to snapshot data of cellular states. This loss of temporal information presents significant challenges in inferring dynamics, as well as causes of cell-to-cell variability. In particular, we are typically unable to separate dynamic variability from within individual systems (“intrinsic noise”) from variability across the population (“extrinsic noise”). Here we mathematically formalise this non-identifiability; but we also use this to identify how new experimental set-ups coupled to statistical noise decomposition can resolve this non-identifiability. For single-cell transcriptomic data we find that systems subject to population variation invariably inflate the apparent degree of burstiness of the underlying process. Such identifiability problems can, in principle, be remedied by dual-reporter assays, which separates total gene expression noise into intrinsic and extrinsic contributions; unfortunately, however, this requires pairs of strictly independent and identical gene reporters to be integrated into the same cell, which is difficult to implement experimentally in most systems. Here we demonstrate mathematically that, in some cases decomposition of transcriptional noise is possible with non-identical and not-necessarily independent reporters. We use our result to show that generic reporters lying in the same biochemical pathways (e.g. mRNA and protein) can replace dual reporters, enabling the noise decomposition to be obtained from only a single gene. Stochastic simulations are used to support our theory, and show that our “pathway-reporter” method compares favourably to the dual-reporter method.

## Introduction

Noise is a fundamental aspect of every cellular process [1]. Frequently it is even of functional importance, for example in driving cell-fate transitions. Sometimes it can afford evolutionary advantages, for example, in the context of bet-hedging strategies. Sometimes, it can be a nuisance, for example, when it makes cellular signal processing more difficult. But noise is nearly ubiquitous at the molecular scale, and its presence has profoundly shaped cellular life. Analysing and understanding the sources of noise, how it is propagated, amplified or attenuated, and how it can be controlled, has therefore become a cornerstone of modern molecular cell biology. Noise arising in gene expression has arguably attracted most of the attention so far (but see e.g. [2] and [3] for the analysis of noise at the signalling level). Generally speaking, gene expression noise is separable into two sources of variability, as pioneered by Swain et al. [4]. Intrinsic noise is generated by the dynamics of the gene expression process itself. The process, however, is often influenced by other external factors, such as the availability of promoters and of RNA polymerase, the influence of long noncoding RNA as a transcriptional regulator [5], as well as differences in the cellular environment. Such sources of variability contribute extrinsic noise, and reflect the variation in gene expression and transcription activity across the cell population. As such, understanding extrinsic noise lies at the heart of understanding cell-population heterogeneity.

So far, elucidating the sources of gene expression noise from transcriptomic measurements alone has proven difficult [6, 7]. The fundamental hindrance lies in the fact that single-cell RNA sequencing, which provides most of the available data, is destructive, so that datasets reflect samples from across a population, rather than samples taken repeatedly from the same cell. As temporal information is lost in such measurements [8], it may be impossible to distinguish temporal variability within individual cells (e.g. burstiness), from ensemble variability across the population (i.e. extrinsic noise). A number of numerical and experimental studies have suggested this confounding effect [9–11], showing that systems with intrinsic noise alone exhibit behaviour that is indistinguishable from systems with both extrinsic and intrinsic noise. This is examined more formally in [12], where we show that the moment scaling behaviour and transcript distribution may be indistinguishable from situations with purely intrinsic noise. The limitations in inferring dynamics from population data are becoming increasingly evident, and a number of studies that seek to address some of these problems have emerged [13, 14].

Here we provide a detailed analysis of the extent to which sources of variability are identifiable from population data. We are able to prove rigorously that it is in general impossible to identify the sources of variability, and consequently, the underlying transcription dynamics, from the observed transcript abundance distribution alone. Systems with intrinsic noise alone can always present identically to similar systems with extrinsic noise. Moreover, we demonstrate that extrinsic noise invariably distorts the apparent degree of burstiness of the underlying system: data which seems “bursty” is not necessarily generated by a bursty process, if there is cell-to-cell variability in e.g. the transcriptional machinery across cells. This has important ramifications for our analysis of experimental data: we cannot assess causes of transcriptional variability, if we do not simultaneously assess cell-to-cell variability in the transcriptional machinery. Our results highlight (in fact prove mathematically) the requirement for additional information, beyond the observed copy number distribution, in order to constrain the space of possible dynamics that could give rise to the same distribution.

This seemingly intractable problem can at least partially be resolved with a brilliantly simple approach: the dual-reporter method [4]. In this approach, noise can be separated into extrinsic and intrinsic components, by observing correlations between the expression of two independent, but identically distributed fluorescent reporter genes. Dual-reporter assays have been employed experimentally to study the noise contributed by both global and gene-specific effects [15–17]. A particular challenge, however, is that dual reporters are rarely identically regulated [17, 18], and are not straightforward to set up in every system. More recently, it has been shown that the independence of dual reporters cannot be guaranteed in some systems [19]. As a result, there have been efforts in developing alternative methods for decomposing noise [18, 20]. Here we develop a widely applicable generalisation (and simplification) of the original dual-reporter approach [4]. We demonstrate that non-identical and not-necessarily independent reporters can provide an analogous noise decomposition. Our result shows that measurements taken from the same biochemical pathway (e.g. mRNA and protein) can replace dual reporters, enabling the noise decomposition to be obtained from a single gene. This completely circumvents the requirement of strictly independent and identically regulated reporter genes. The results obtained from our “pathway-reporter” method are also borne out by stochastic simulations, and compare favourably with the dual-reporter method. In the case of constitutive expression, the results obtained from our decomposition are identical to those obtained from dual reporters. For bursty systems, we show that our approach provides a satisfactorily close approximation, except in extreme cases.

## Modelling Intrinsic and Extrinsic Noise

A simple model for stochastic mRNA dynamics is the Telegraph model: a two-state model describing promoter switching, transcription, and mRNA degradation. In this model, all parameters are fixed, and gene expression variability arises due to the inherent stochasticity of the transcription process. As discussed above, this process will often be influenced by extrinsic sources of variability, and so modifications to account for this additional source of variability are required.

### The Telegraph Model

The Telegraph model was first introduced in [21], and since then has been widely employed in the literature to model bursty gene expression in eukaryotic cells [22–25]. In this model, the gene switches probabilistically between an active state and an inactive state, at rates *λ* (on-rate) and *µ* (off-rate), respectively. In the active state, mRNAs are synthesized according to a Poisson process with rate *K*, while in the inactive state, transcription does either not occur, or possibly occurs at some lower Poisson rate, *K*_0_ *≪ K*. Degradation of RNA molecules occurs independently of the gene promoter state at rate *δ*. Fig. 1A shows a schematic of the Telegraph model. Throughout the discussion here, we will rescale all parameters of the Telegraph model by the mRNA degradation rate, so that *δ* = 1. The steady-state distribution for the mRNA copy number can be explicitly calculated as [26],

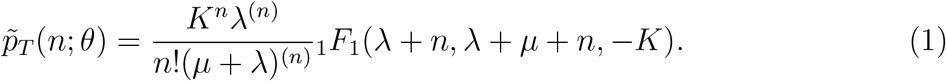

**Figure 1:**
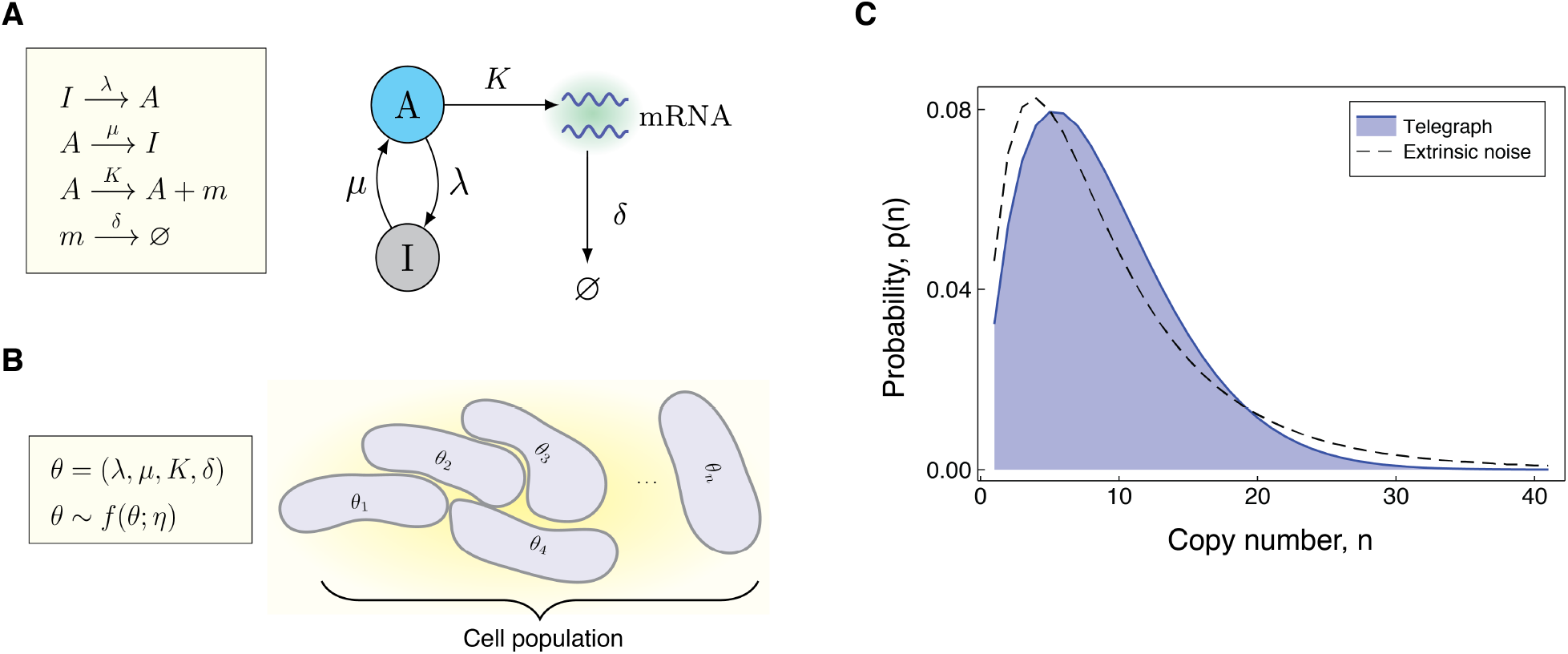
Modelling the effects of both intrinsic and extrinsic noise. (A) A schematic of the Telegraph process, with nodes *A* (active) and *I* (inactive) representing the state of the gene. Transitions between the states *A* and *I* occur stochastically at rates *µ* and *λ*, respectively. The parameter *K* is the mRNA transcription rate, and *δ* is the degradation rate. (B) The compound model incorporates extrinsic noise by assuming that parameters *θ* of the Telegraph model vary across an ensemble of cells, according to some probability distribution *f* (*θ*; *η*). (C) Variation in the parameters across the cell population leads to greater variability in the mRNA copy number distribution.

Here *θ* denotes the parameter vector (*µ, λ, K, δ*), the function _1_*F*_1_ is the confluent hypergeometric function [27], and, for real number *x* and positive integer *n*, the notation *x*^(*n*)^ abbreviates the rising factorial of *x* (also known as the Pochhammer function). Throughout, we will refer to the probability mass function 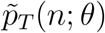 as the Telegraph distribution with parameters *θ*. Constitutive gene expression is just a limiting case of the Telegraph model, which arises when the off-rate *µ* is 0, so that the gene remains permanently in the active state. In this case, the Telegraph distribution simplifies to a Poisson distribution with rate *K*; Pois(*K*).

At the opposite extreme is instantaneously bursty gene expression in which mRNA are produced in very short bursts with potentially prolonged periods of inactivity in between. This mode of gene expression has been frequently reported experimentally, particularly in mammalian genes [17, 22, 24, 25]. Transcriptional bursting may be treated as a limit of the Telegraph model, where the off-rate, *µ*, has tended toward infinity, while the on-rate *λ* remains fixed. In this limit, it can be shown [9, 12] that the Telegraph distribution converges to the negative binomial distribution NegBin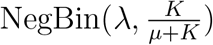.

### The Compound Distribution

We can elegantly account for random cell-to-cell variation across a population by way of a compound distribution [28]

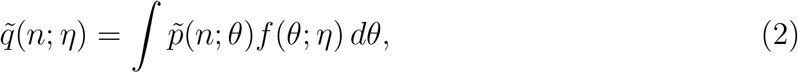

where 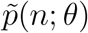 is the stationary probability distribution of a system with fixed parameters *θ* and *f* (*θ*; *η*) denotes the multivariate distribution for *θ* with hyperparameters *η*. Often we will take 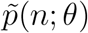 to be the stationary probability distribution of the Telegraph model ((1)), and refer to (2) as the compound Telegraph distribution. Sometimes 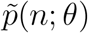 will be the Poisson distribution or the negative binomial distribution, depending on the underlying mode of gene activity.

The compound distribution is valid in the case of ensemble heterogeneity, that is, when parameter values differ between individual cells according to the distribution *f* (*θ*; *η*), but remain constant over time [2]. This model is also a valid approximation for individual cells with dynamic parameters, provided these change sufficiently slower than the transcriptional dynamics [29]. Fig. 1B gives a pictorial representation of the compound distribution.

In general, the compound Telegraph distribution 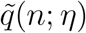 will be more dispersed than a Telegraph distribution to account for the uncertainty in the parameters; see Fig. 1C. Such dispersion is widely observed experimentally, and as demonstrated in [12], reflects the presence of extrinsic noise.

### Identifiability Considerations

Decoupling the effects of extrinsic noise from experimental measurements has been notoriously challenging. In the context of (2), the distribution *f* (*θ*; *η*) reflects the population heterogeneity, but experimental data provides samples only of 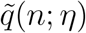. How much can we deduce of the underlying dynamics (that is, 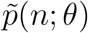), and the population heterogeneity (*f* (*θ*; *η*)), from measurements of transcripts from across the cell population 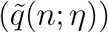

Of course, even though we may be able to infer the parameter *η* from experimental data, the expression 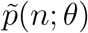 is really a family of distributions, parameterised by *θ*. This presents two fundamental challenges. The first is the possibility that there are different families of distributions 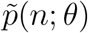 that can yield the same compound distribution, 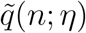, but generated by different mechanisms, i.e. noise distributions, *f* (*θ*; *η*). The second, perhaps more subtle, challenge is that, even for a fixed family of distributions 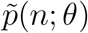 it may be possible that different choices of the noise distribution *f* (*θ*; *η*) could still yield the same compound distribution 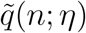. In these situations, we cannot hope to infer a unique solution for the noise distribution. This belongs to the important class of identifiability problems [30]; it has important ramifications for the interpretability of parameter estimates obtained from experimental data [31]. Indeed, if two or more model parameterisations are observationally equivalent (in this case, in the form of the transcript abundance distribution 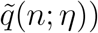, then not only does this cast doubt upon the suitability of the model to represent (and subsequently predict) the system, but it also obstructs our ability to infer mechanistic insight from experimental data.

An example of the first identifiability problem arises from a well-known example of a compound distribution, (2): when *f* (*θ*; *η*) is a gamma distribution and 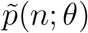 is a Poisson distribution, corresponding to constitutive gene expression, the resulting compound distribution 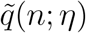 is a negative binomial distribution [32]. But this is the same distribution as that arising from instantaneously bursty gene expression [12]. Such identifiability instances may be circumvented if there is confidence in the basic mode of gene activity, i.e., if there is reason to believe that a gene is not constitutively active, for example. We find, however, that there are numerous instances that can present insurmountable identifiability problems.

### Bursty Gene Expression

We first observe that any Telegraph distribution with fixed parameters can be identically obtained from a Telegraph distribution with parameter variation. As shown in the supplementary material, any Telegraph distribution 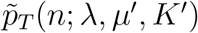 can be written as,

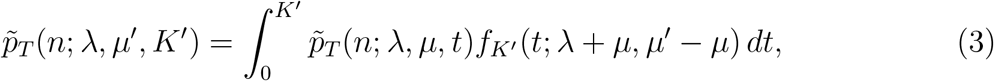

where *µ < µ*^*′*^ and *f*_*K′*_(*t*; *λ* + *µ, µ*^*′*^ *− µ*) is the probability density function for a scaled beta distribution Beta_*K*_(*λ* + *µ, µ*^*′*^ *− µ*) with support [0, *K*^*′*^]. Thus, any Telegraph distribution can be obtained by varying the transcription rate parameter on a narrower Telegraph distribution (i.e., with a smaller off-rate) according to a scaled beta distribution. Fig. 2A (top panel) compares the representation obtained in (3) with the corresponding fixed-parameter Telegraph distribution for two different sets of parameters. When *µ* = 0 the representation given in (3) simplifies to the well-known Poisson representation of the Telegraph distribution in terms of the scaled beta distribution [33].

**Figure 2:**
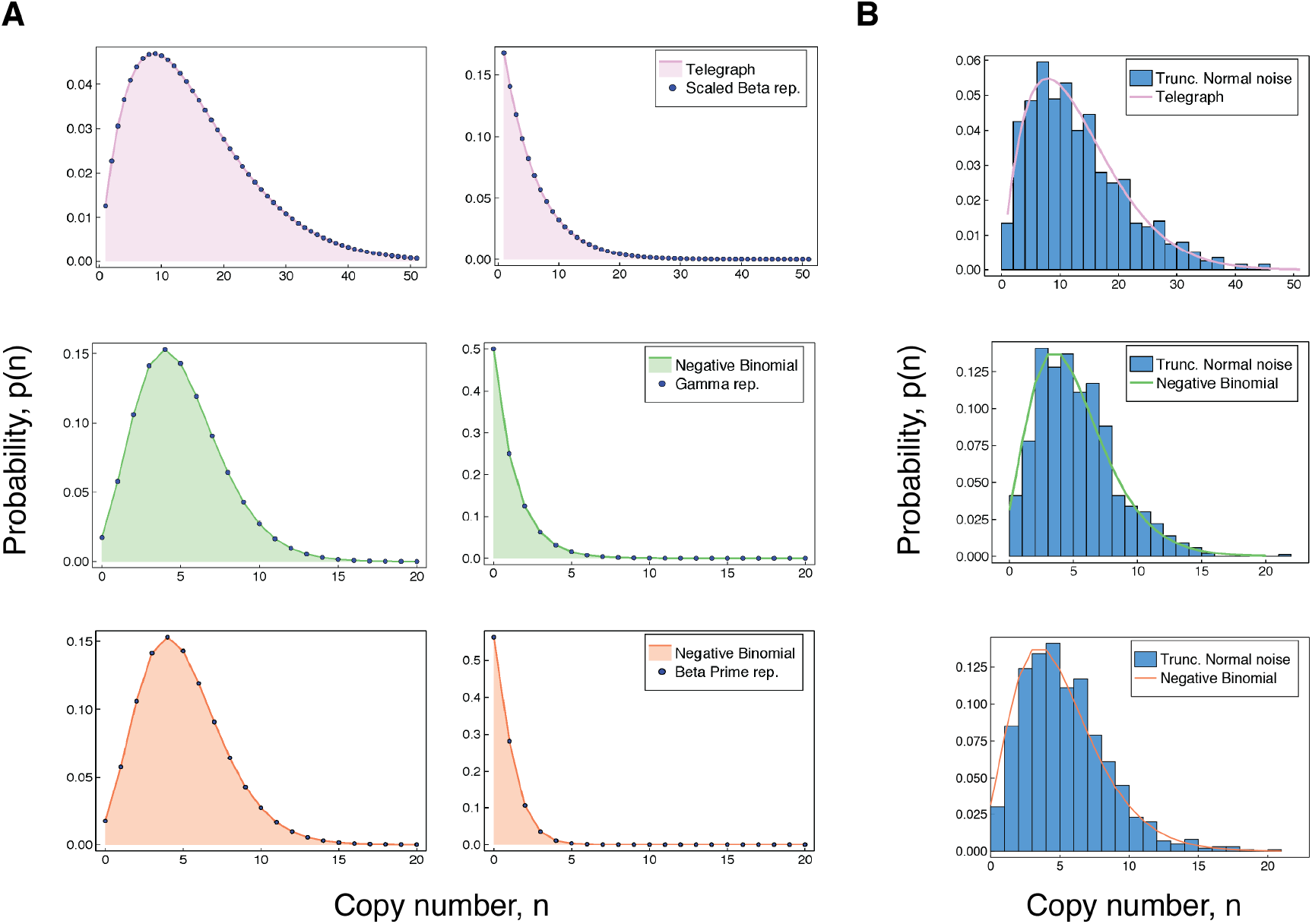
(A) Accuracy of our representations for the Telegraph and negative binomial distribution. For each of the results in (3)-(5), we compare the (fixed-parameter) Telegraph and negative binomial distributions with their respective compound representations for two different sets of parameter values. The top panel (shown in pink) is shows comparisons for (3). Referring to Table 1, the parameter values for the top panel are (left) *λ* = 2, *µ*^*′*^ = 12, *K*^*′*^ = 100, *µ* = 3, and *K ∼* Beta_*K′*_(5, 9), and (right) *λ* = 1, *µ*^*′*^ = 20, *K*^*′*^ = 100, *µ* = 2 and *K ∼* Beta_*K′*_ (3, 18). The middle panel (green) gives comparisons for (4), with parameter values (left) *λ* = 10, *β* = 2, *µ* = 2 and *K ∼* Gamma(12, 2) and (right) *λ* = 1, *β* = 1, *µ* = 2 and *K ∼* Gamma(3, 1). The bottom panel (coral) gives comparisons for (5). The parameter values (left) are *λ*^*′*^ = 10, *λ* = 15 and *c* = 2 and (right) are *λ*^*′*^ = 2, *λ* = 5 and *c* = 3. (B) The top figure compares a Telegraph(2, 4, 60) distribution with samples from a compound Telegraph distribution with normal noise Norm(37, 10) on the transcription rate parameter. The middle figure compares a NegBin(5, 0.5) with samples from a compound Telegraph distribution with normal noise Norm(5.5, 2.3) on the transcription rate parameter. The bottom figure compares a NegBin(5, 1) distribution with samples from a compound negative binomial distribution with normal noise Norm(2.3, 0.6) on the burst intensity parameter.

### Instantaneously Bursty Gene Expression

The previous result extends to instantaneously bursty systems. The copy number distribution of an instantaneously bursty system can be obtained from both bursty and instantaneously bursty dynamics, provided that there is appropriate parameter variation. The supplementary material contains the relevant derivations. In the following, we let 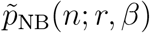 denote the probability mass function of a 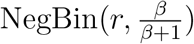 distribution, where *β >* 0. Then for any negative binomial distribution of the form 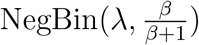 we have,

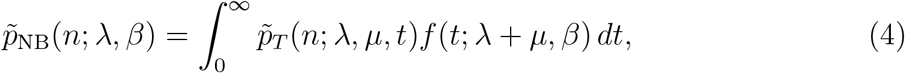

where *f* (*t*; *λ* + *µ, β*) is the probability density function of a Gamma(*λ* + *µ, β*) distribution. This result generalises the aforementioned classic representation of the negative binomial distribution [32], which corresponds to the particular case of *µ* = 0. In Fig. 2A (middle panel), we compare the representation obtained in (4) with the corresponding fixed-parameter negative binomial distribution for two different sets of parameters.

**Table 1:** Summary of the non-identifiability results. The results given in lines 1, 3 and 5 are our contributions, while the remaining representations (lines 2 and 4) are known and can be obtained as special cases of our results. Note that here we use Tele(*λ, µ, K*) to denote a Telegraph distribution with parameters *λ, µ, K*. In lines 3 and 4, the parameter *β >* 0 can be chosen freely and determines the mean burst intensity in the resulting compound system. In line 5 the parameters *b, θ >* 0 are again mean burst intensities, and *b* can be chosen freely in the determination of the distribution of *θ*.

| Copy no. distribution $\tilde{q}(n; \eta)$ | Underlying distribution $\tilde{p}(n; \theta)$ | Noise distribution $f(\theta; \eta)$ |
| --- | --- | --- |
| Tele($\lambda, \mu', K'$) | Tele($\lambda, \mu, K$) | $K \sim \text{Beta}_{K'}(\lambda + \mu, \mu' - \mu)$ |
| Tele($\lambda, \mu', K'$) | Pois($K$) | $K \sim \text{Beta}_{K'}(\lambda, \mu')$ |
| NegBin($\lambda, \frac{\beta}{\beta+1}$) | Tele($\lambda, \mu, K$) | $K \sim \text{Gamma}(\lambda + \mu, \beta)$ |
| NegBin($\lambda, \frac{\beta}{\beta+1}$) | Pois($K$) | $K \sim \text{Gamma}(\lambda, \beta)$ |
| NegBin($\lambda', \frac{b}{b+1}$) | NegBin($\lambda, \frac{\theta}{\theta+1}$) | $\theta \sim \text{BetaPrime}_b(\lambda - \lambda', \lambda')$ |

We also have the following representation for a negative binomial distribution in terms of a scaled beta prime distribution,

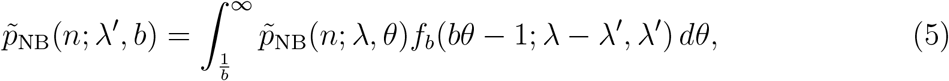

where *f*_*b*_(*bθ−*1; *λ−λ*^*′*^, *λ*^*′*^) is the probability mass function of a scaled beta prime BetaPrime_*b*_(*λ−λ*^*′*^, *λ*) distribution, where *b >* 0 and *λ > λ*^*′*^. This equivalently corresponds to scaled beta noise Beta_*b*_(*λ−λ*^*′*^, *λ*^*′*^) on the inverse of the expected burst intensity. Thus, the distribution of any instantaneously bursty system with mean burst intensity *b* can be obtained from one with greater burst frequency, by varying the mean burst intensity *θ* according to a shifted beta prime distribution. Fig. 2A (bottom panel), compares the representation obtained in (5) with the associated fixed-parameter negative binomial distribution for two different sets of parameters.

### An Exception: Constitutive Expression

It has long been known [34] that a compound Poisson distribution uniquely determines the compounding distribution. In the context of (2), this means the full extrinsic noise distribution *f* (*θ, η*) is identifiable from 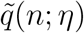. As we demonstrate in related work [35], it is therefore possible to extract the extrinsic noise distribution, *f* (*θ, η*), from transcript copy number measurements.

### Implications for Parameter Inference

Estimates of kinetic parameters from experimental data suggest that gene expression is often either bursty or instantaneously bursty (i.e., *µ ≫ λ*). In turn, the assumption that gene-inactivation events occur far more frequently than gene-activation events is often used to derive other models of stochastic mRNA dynamics [36–38]. The representations given in (3)–(5), however, show that both estimating parameters and the underlying dynamics from the form of the copy number distribution alone can be misleading. Noise on the transcription rate will invariably produce copy number data that is suggestive of a more bursty model. To illustrate this, consider an example in which the underlying process is a (mildly) bursty Telegraph system with distribution 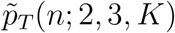. Now assume that noise on the transcription rate parameter *K* follows the scaled Beta distribution on the interval [0, 100] with *α* = *λ* + *µ* = 2 + 3 = 5 and *β* = *µ*^*′*^ *− µ* = 12 *−* 3 = 9. The shape of this noise distribution closely resembles a slightly skewed Gaussian distribution, with the majority of transcription rates between around 10 and 60. This noise on the transcription rate *K* within the Telegraph system 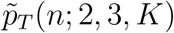 will present identically to the significantly burstier system 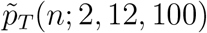.

It is of practical importance to recognise that, while the non-identifiability results (summarised in Table 1) are dependent on specific noise distributions, for practical purposes any similar distribution will produce a similar effect. To demonstrate this, we replace the various noise distributions required for the representations in (3)–(5), with suitable normal distributions truncated at 0. In each case, we sample 1,000 data points from the corresponding compound distribution, and compared this with the associated fixed-parameter copy number distribution. The results are shown in Fig. 2B. The truncated normal distribution is not chosen on the basis of biological relevance, but rather to demonstrate that even a symmetric noise distribution (except for truncation at 0) produces qualitatively similar results to the distributions used in the precise non-identifiability results. In every case, the effect of a unimodal noise distribution on the transcription rate or burst intensity parameter is to produce copy number distributions that are generally consistent with systems that appear burstier.

Finally, we note that our results explain previous empirical observations that static measurements of mRNA are not always sufficient to infer the underlying dynamics of gene activity. Skinner et al. [13] address some of these limitations by quantifying both nascent and mature mRNA in individual cells, as well incorporating cell-cycle effects into their analysis of two mammalian genes. A more developed treatment of model identifiability is given in [14], where it shown how a stochastic model incorporating the downstream processing of mRNA can be used to distinguish a particular instance of non-identifiability. More specifically, the authors consider the non-identifiability problem noted in [12], arising from the Gamma-Poisson compound representation of the negative binomial distribution; a particular case of (4) above. Despite identical distributions at the nascent level, the marginal distributions of the processed (mature) mRNA are found to be substantially different. It is likely that a similar analysis will be valuable in the context of the other identifiability problems we have given in Table 1. In the next section, however, we take a more general approach to resolving non-identifibiality and exploit the properties of complex gene expression dynamics to determine not only the presence of extrinsic noise, but also estimate its magnitude.

## Resolving Non-identifiability

The results of the previous section show that additional information, beyond the observed copy number distribution, is required to constrain the space of possible dynamics that could give rise to the same distribution. One way to narrow this space of possibilities, is to determine the intrinsic and extrinsic contributions to the total variation in the system.

### The Dual-Reporter Method

The total gene expression noise, as measured by the squared coefficient of variation *η*^2^, can be decomposed exactly into a sum of intrinsic and extrinsic noise contributions [4]. The decomposition applies to dynamic noise [39], and generalises to higher moments in [40].

As shown in [39], the noise decomposition is equivalent to the normalised Law of Total Variance [41]. Indeed, if *X* is the random variable denoting the number of molecules of a certain species (eg. mRNA or protein) in a given cell, then we can decompose the total noise by conditioning *X* on the state **Z** = (*Z*_1_, …, *Z*_*n*_) of the environmental variables *Z*_1_,…,*Z*_*n*_,

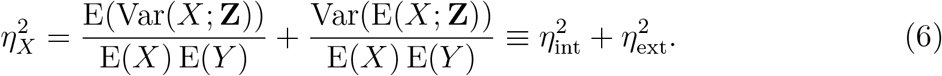

It has been shown [4, 39] that if *X*_1_ and *X*_2_ are random variables denoting the expression levels of independent and identically distributed gene reporters, then the extrinsic noise contribution 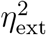 in (6) can be identified by the normalised covariance between *X*_1_ and *X*_2_,

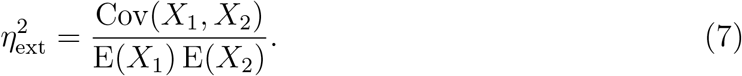

### Decomposing Noise with Non-Identical Reporters

The dual-reporter method requires distinguishable measurements of transcripts or proteins from two independent and identically distributed reporter genes integrated into the same cell. In practice, however, dual reporters rarely have identical dynamics, which is widely considered to be a significant challenge to interpreting experimental results [18]. We show that, under certain conditions, the decomposition in (6) can alternatively be obtained from non-identically distributed and not-necessarily independent reporters.

Our result relies on the observation that the covariance of any two variables can be decomposed into the expectation of a conditional covariance and the covariance of two conditional expectations (the Law of Total Covariance [41]). If *X* and *Y* denote, for example, the numbers of molecules of two chemical species (eg. mRNA and protein) in a given cell, then the covariance of *X* and *Y* can be decomposed by conditioning on the state **Z** = (*Z*_1_, …, *Z*_*n*_) of the environmental variables *Z*_1_,…,*Z*_*n*_,

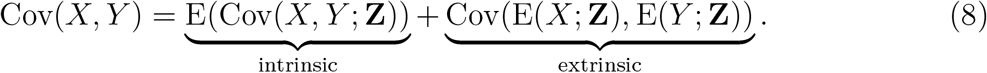

We will see that in many cases of interest the random variable E(*X*; **Z**) (as a function of **Z**) splits across common variables with E(*Y*; **Z**). By this we mean that E(*X*; **Z**) = *f* (**Z**_*X*_)*h*_*X*_(**Z**^*′*^) and E(*Y*; **Z**) = *g*(**Z**_*Y*_)*h*_*Y*_ (**Z**^*′*^), where **Z**_*X*_ are the variables of **Z** that appear in E(*X*; **Z**) but not in E(*Y*; **Z**), and dually, **Z**_*Y*_ are those in E(*Y*; **Z**) that are not in E(*X*; **Z**). The variables **Z**^*′*^ are those variables from **Z** not in **Z**_*X*_∪**Z**_*Y*_. In these cases, the component of Cov(*X, Y*) that is contributed by the variation in **Z** (the extrinsic component) may be written as the covariance of the functions *h*_*X*_(**Z**^*′*^) and *h*_*Y*_ (**Z**^*′*^). Conveniently, in the cases of interest here, the two functions *h*_*X*_ and *h*_*Y*_ coincide, and this is the form we use in the following decomposition principle. The supplementary material contains the proof of this result.

### The Noise Decomposition Principle (NDP)

Assume that there are measurable functions *f, g* and *h* such that E(*X*; **Z**) and E(*Y*; **Z**) split across common variables by way of E(*X*; **Z**) = *f* (**Z**_*X*_)*h*(**Z**^*′*^) and E(*Y*; **Z**) = *g*(**Z**_*Y*_)*h*(**Z**^*′*^). Then, provided that the variables *Z*_1_, …, *Z*_*m*_ are mutually independent, the normalised covariance of E(*X*; **Z**) and E(*Y*; **Z**) will identify the total noise on *h*(**Z**^*′*^) (i.e., 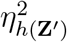).

As we show in the next section, there are many situations where the random variable E(*X*; **Z**) is precisely the common part of E(*Y*; **Z**) and E(*X*; **Z**) (i.e., *h*(**Z**^*′*^) = E(*X*; **Z**)), and the normalised intrinsic contribution to the covariance is either zero or negligible. In these cases, the normalised covariance of *X* and *Y* will identify precisely the extrinsic noise contribution 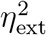 to the total noise 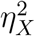. To see this, consider the situation where E(*Y*; **Z**) = *f* (**Z**_*Y*_) E(*X*; **Z**). Then provided *f* (**Z**_*Y*_) and E(*X*; **Z**) are independent random variables, the extrinsic contribution to the covariance of *X* and *Y* is given by,

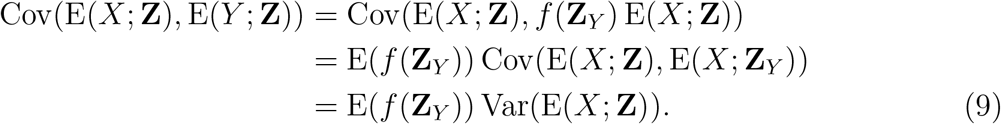

If the normalised intrinsic contribution to the covariance is either zero or is negligible, it follows from (8) that

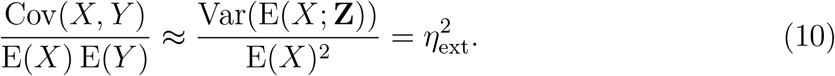

Thus, under certain conditions, measuring the covariance between two non-identically distributed and not-necessarily independent reporters can replace dual reporters.

### The Pathway-Reporter Method

We show that for some reporters *X* and *Y* belonging to the same biochemical pathway, the covariance of *X* and *Y* continues to identify the extrinsic, and subsequently intrinsic, noise contributions to the total noise. The basis of the pathway-reporter method depends on the emergent covariances between the various species (for eg. nascent/mature mRNA and protein) in the gene expression pathway. Qualitatively, this effect can be seen in Figure 3, which compares simulated sample distributions of a simple four-stage model of gene transcription (refer to the model **M**_4_ below) in the case of moderate extrinsic noise to the case with no extrinsic noise. The plots are the bivariate distributions for nascent-mature, nascent-protein and mature-protein levels, respectively. This will be made more precise below, where we find that it is possible to extract quantitative information about the extrinsic noise distribution itself from this data.

**Figure 3:**
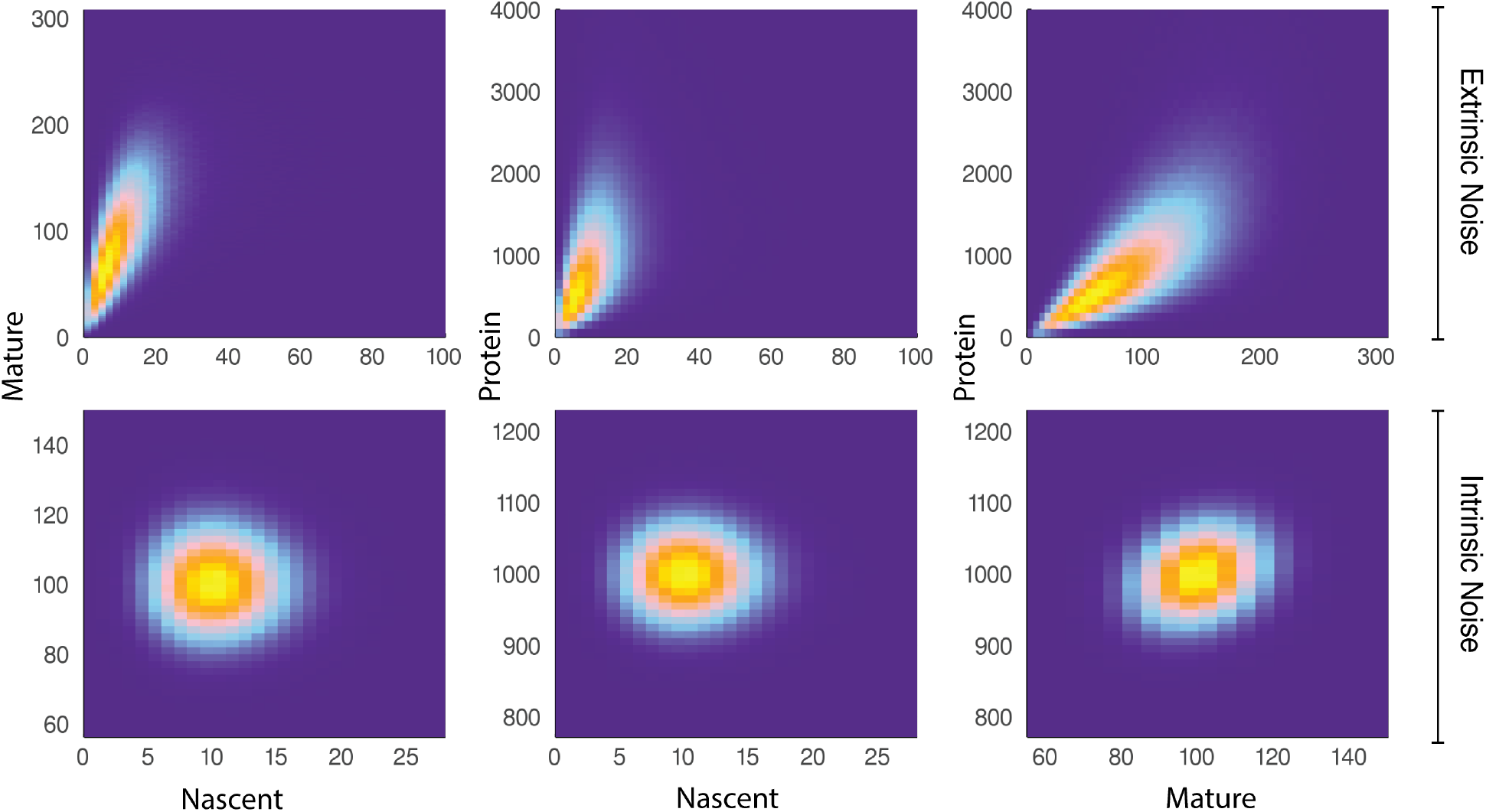
A comparison of joint distributions in the case of moderate extrinsic noise and no extrinsic noise. The plots are generated from a three-stage model of gene transcription, incorporating the production of nascent mRNA, mature mRNA and protein. Details of the model can be found in Fig. 4 (model **M**_4_) and the associated text. The top panel shows nascent-mature, nascent-protein and mature-protein joint distributions in the case of extrinsic noise, while the bottom panel displays the corresponding plots in the case of no extrinsic noise. Extrinsic noise produces a visibly more correlated joint distribution, which forms the basis of the pathway-reporter method.

Throughout this section, we assume that extrinsic noise sources act independently i.e., the environmental variables *Z*_1_, …, *Z*_*n*_ of **Z** are mutually independent. Additionally, our modelling focuses only on a single gene copy, though the same analysis applies to multiple but indistinguishable gene copies; we refer to the supplementary material for more details.

### Measuring Noise from a Constitutive Gene

We consider first the simplest case where the underlying process is constitutive. We begin with a stochastic model of mRNA maturation, which we denote by **M**_1_; Fig. 4A (top left) gives a schematic of the constitutive model. In this model, the gene continuously produces nascent mRNA according to a Poisson process at constant rate *K*_*N*_, which are subsequently spliced into mature mRNA according at rate *K*_*M*_. Degradation of mature mRNA occurs as a first-order Poisson process with rate *δ*_*M*_. The model **M**_1_, together with its extensions, has been considered in a number of recent studies [14, 37, 42–44], and has a known solution for the stationary joint probability distribution [45] given by,

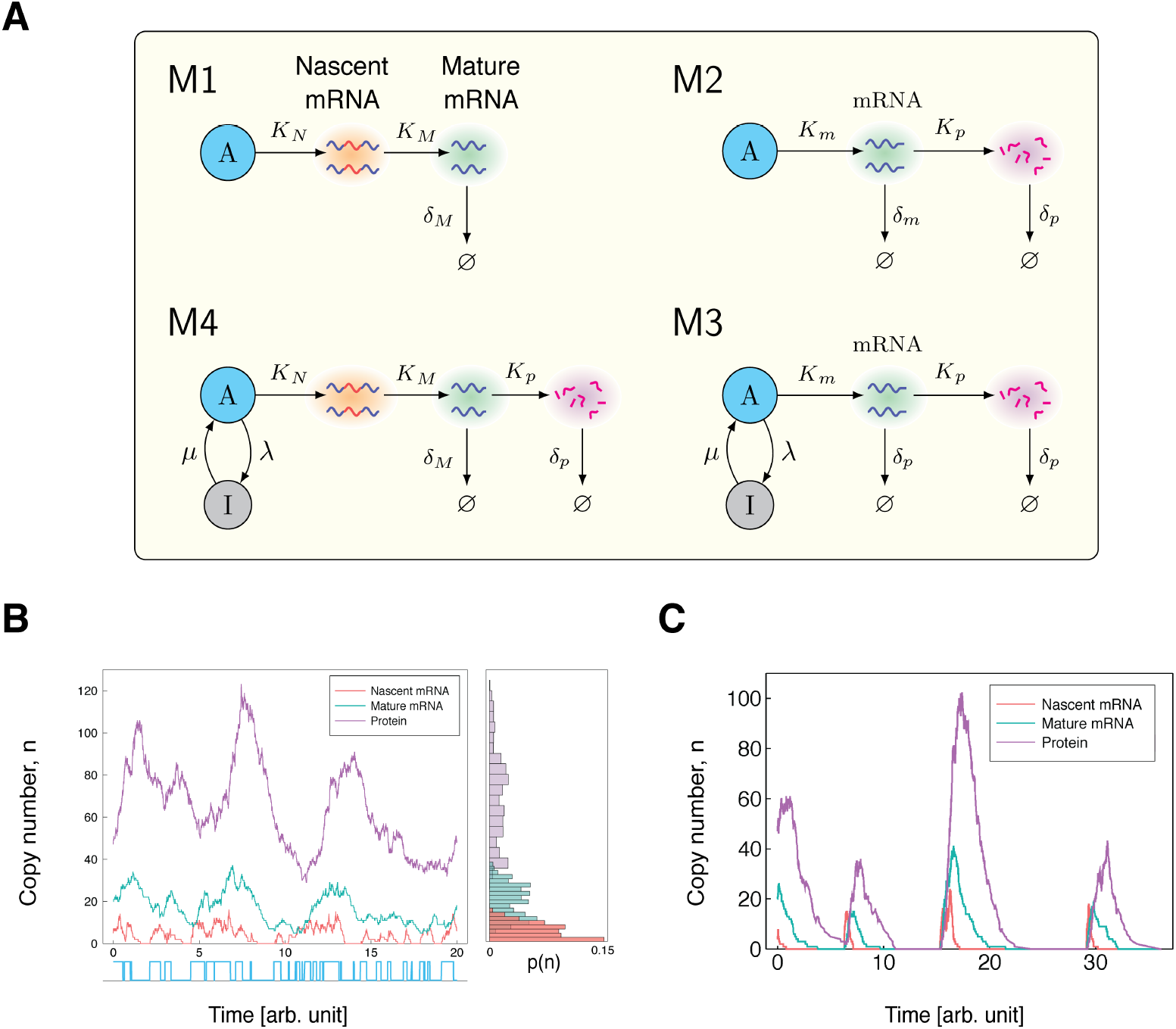

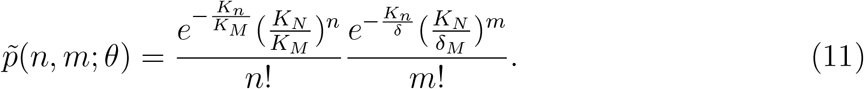

Here *n* is the number of nascent mRNA, *m* is the number of mature mRNA, and *θ* = (*K*_*N*_, *K*_*M*_, *δ*_*M*_). We let *X*_*N*_ denote the number of nascent mRNA, *X*_*M*_ the number of mature mRNA produced from the same constitutive gene, and **Z** = (*K*_*N*_, *K*_*M*_, *δ*_*M*_). To simplify notation, we abbreviate the variables in 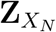 as **Z**_*N*_, and similarly for 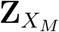. It follows immediately from (11) that *X*_*N*_ and *X*_*M*_ are independent conditional on **Z**, and so the intrinsic contribution to the covariance of *X*_*N*_ and *X*_*M*_ (the first term of (8)) is 0. It is also easy to see from (11) that E(*X*_*N*_; **Z**) = *f* (**Z**_*N*_)*K*_*N*_ and E(*X*_*M*_; **Z**) = *g*(**Z**_*M*_)*K*_*N*_, where 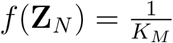 and 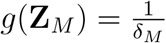. Since the extrinsic noise sources are assumed to act independently, it follows that the Noise Decomposition Principle (NDP) of the previous section holds. We then have that 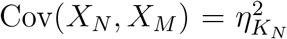, where 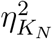 is the total noise on the transcription rate parameter *K*_*N*_. Thus, measuring Cov(*X*_*N*_, *X*_*M*_) can replace dual reporters to decompose the gene expression noise at the transcriptional level.

To support our mathematical results, we simulate the model **M**_1_ subject to parameter variation using the stochastic simulation algorithm (SSA). Table 2 compares the extrinsic noise contributions found from various simulations with the corresponding theoretical values. In each simulation, the degradation rate *δ*_*m*_ is fixed at 1, with the other parameters scales accordingly. The maturation rate *K*_*M*_ is sampled according to a Gamma(8, 0.0125) distribution, which has coefficient of variation 0.125. We consider different noise distributions on *K*_*N*_, producing a range of noise strengths. Our theory predicts that pathway-reporters will identify the total noise on *K*_*N*_. Overall, we observe an excellent agreement between the results obtained by the pathway-reporter method, the dual-reporter method and the theoretical noise. There is consistently slightly more variation in the pathway-reporter results compared with the dual-reporter results.

**Table 2:** A comparison of the pathway-reporter method and the dual-reporter method for constitutive expression under the model **M**_1_. Here PR gives the results of the nascent and mature pathway reporters, while DR (Mat) gives the results of dual reporters calculated from the mature mRNA. We considered noise on both the transcription rate (*K*_*N*_) and the maturation rate (*K*_*M*_). The decay rate is fixed at one, with the other parameters scaled accordingly. In each case, the maturation rate *K*_*M*_ is varied according to a Gamma(8, 1.25) distribution, which has coefficient of variation 0.125. The values given are the average of 100 simulations, each calculated from 500 copy number samples, and the errors are ± one standard deviation. Our theory predicts that pathway-reporters will identify the noise on the nascent transcription rate *K*_N_ 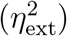). The noise distribution parameters are chosen to produce an average nascent mRNA copy number of approximately 5 and an average mature mRNA copy number of approximately 50.

| Theory |  | Simulation |  |
| --- | --- | --- | --- |
| $\eta_{\text{ext}}^2$ | Noise ($K_N$) | PR | DR (Mat) |
| 0.00 | $K_N = 50$ | $0.00 \pm 0.01$ | $0.00 \pm 0.00$ |
| 0.10 | Beta<sub>133.3</sub>(6, 10.5) | $0.10 \pm 0.01$ | $0.10 \pm 0.01$ |
| 0.20 | Gamma(5, 10) | $0.20 \pm 0.02$ | $0.20 \pm 0.01$ |
| 0.50 | Beta<sub>300</sub>(1.5, 7.5) | $0.49 \pm 0.04$ | $0.50 \pm 0.03$ |

To explore the pathway-reporter method more widely, we consider 60 different parameter combinations to produce a range of mean copy numbers consistent with those reported experimentally. We also consider different noise distributions taken from the scaled Beta distribution family in order to produce a range of noise strengths; see the supplementary dataset S1. The pathway-reporter method performa favourably compared to the dual-reporter method calculated from mature mRNA, and consistently outperforms the dual-reporter method on nascent mRNA. Next, we consider the simplest stochastic model of gene expression that includes both mRNA and protein dynamics: the well-known “two-stage model” of gene expression, which, together with its three-stage extension to include promoter switching has been widely studied [16, 17, 46–51]. We denote this model by **M**_2_; see Fig. 4A (top right) for a schematic of this model. In this model, mRNA are synthesized according a Poisson process at rate *K*_*m*_, which are then later translated into protein at rate *K*_*p*_. Degradation of mRNA and protein occur as first-order Poisson processes with rates *δ*_*m*_ and *δ*_*p*_, respectively. If *X*_*m*_ denotes the number of mRNA, *X*_*p*_ the number of proteins produced from the same constitutive gene, and if **Z** = (*K*_*m*_, *K*_*p*_, *δ*_*m*_, *δ*_*p*_), then the stationary means and covariance are given by [46, 48]:

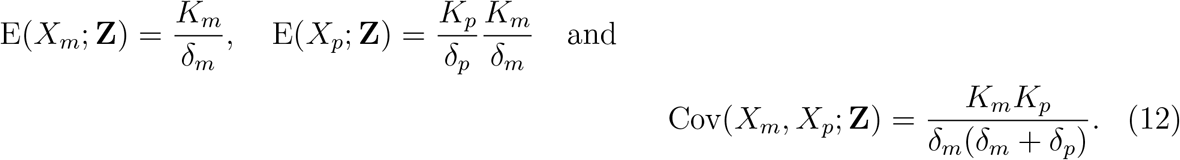

It is easily verified that E(*X*_*p*_; **Z**) = *f* (**Z**_*p*_) E(*X*_*m*_; **Z**), where 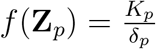. Thus, it follows from the NDP that the normalised contribution of Cov(*X*_*m*_, *X*_*p*_) contributed by **Z** will identify the extrinsic noise contribution to the total noise on *X*_*m*_. Now, if we assume that *δ*_*m*_ is fixed across the cell-population, and all parameters are scaled so that *δ*_*m*_ = 1, we have the following expression for the intrinsic contribution to the covariance of *X*_*m*_ and *X*_*p*_ (refer to the supplementary material for details),

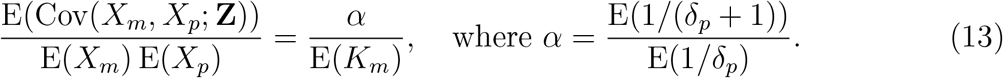

Since mRNA tends to be less stable than protein, we have *δ*_*p*_ *<* 1, and often *δ*_*p*_ *≪* 1 [52, 53]. So, we can expect *α* ≪ 1. Further, for many genes we can expect the number of mRNA per cell (*K*_*m*_) to be in the order of tens, so 1*/E*(*K*_*m*_) *<* 1. It follows that E(Cov(*X*_*m*_, *X*_*p*_; **Z**)) ≪ 1, so that Cov(*X*_*m*_, *X*_*p*_) will closely approximate the extrinsic noise at the transcriptional level.

We tested our theory using stochastic simulations of the model **M**_2_ subject to parameter variation. Table 3 gives a comparison of the results of the mRNA-protein reporters and dual reporters. In each case, we varied *K*_*p*_ according to a Gamma(5, 0.4) distribution and *δ*_*p*_ according to a Gamma(8, 0.125) distribution; the corresponding noise strengths are 0.20 and 0.125, respectively. We consider different noise distributions on *K*_*m*_, which produce a range of noise strengths, and the noise distribution parameters are selected to produce a mean mRNA of approximately 50 and a mean number of approximately 1000 proteins in each simulation. As our theory predicts, the mRNA-protein reporters identify the extrinsic noise contribution to the total noise on *X*_*m*_. Again, we can see from Table 3 that there is excellent agreement between the results of the pathway reporters and the dual reporters, with slightly more variation in the pathway-reporter results. A larger exploration of the parameter space reveals similar results; these are provided in the supplementary dataset S1. Thus, despite mRNA and protein numbers not being strictly independent, they can, for practical purposes, replace dual reporters to decompose the noise at the transcriptional level.

**Table 3:**
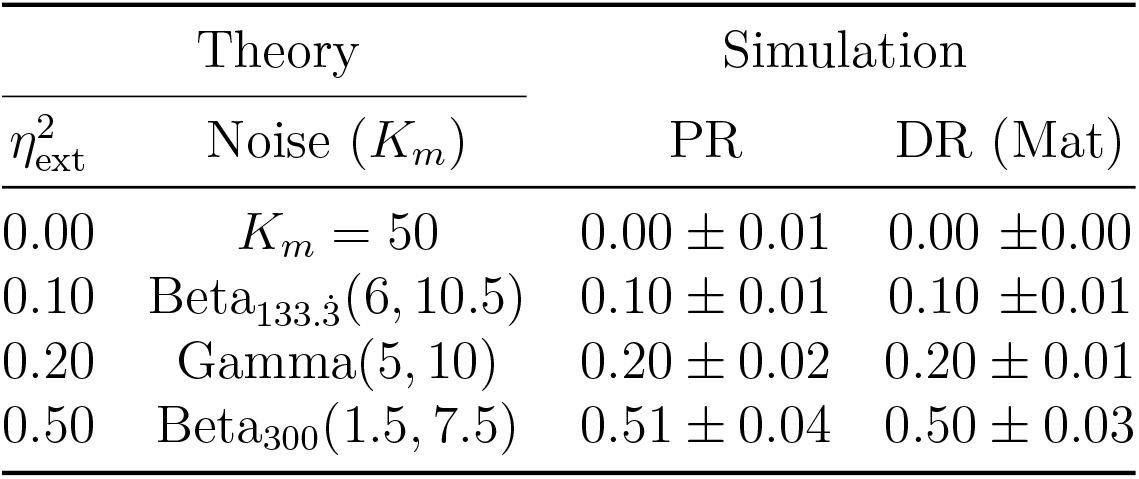
A comparison of the pathway-reporter method and the dual-reporter method for constitutive expression under the model **M**_2_. Here PR gives the results of the mRNAprotien pathway reporters, while DR (Mat) gives the results of dual reporters calculated from the mature mRNA. We considered noise on the transcription rate (*K*_*m*_), the protein synthesis rate (*K*_*p*_), and the protein decay rate (*δ*_*p*_). The mRNA decay rate is fixed at one. In each case, we varied *K*_*p*_ according to a Gamma(5, 0.4) distribution and *δ*_*p*_ according to a Gamma(8, 0.125) distribution; the corresponding noise strengths are 0.20 and 0.125, respectively. We considered different noise distributions on *K*_*m*_, which produce a range of noise strengths. The noise distribution parameters are selected to produce a mean mRNA of approximately 50 and a mean number of approximately 1000 proteins in each simulation. The values given are the average of 100 simulations, each calculated from 500 copy number samples, and the errors are ± one standard deviation. As our theory predicts, the mRNA-protein reporters identify the noise on the transcription rate parameter *K*_*m*_ 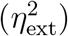.

| $\eta_{\text{ext}}^2$ | Theory | Simulation | |
| --- | --- | --- | --- |
| | Noise ($K_m$) | PR | DR (Mat) |
| 0.00 | $K_m = 50$ | $0.00 \pm 0.01$ | $0.00 \pm 0.00$ |
| 0.10 | Beta<sub>133.3</sub>(6, 10.5) | $0.10 \pm 0.01$ | $0.10 \pm 0.01$ |
| 0.20 | Gamma(5, 10) | $0.20 \pm 0.02$ | $0.20 \pm 0.01$ |
| 0.50 | Beta<sub>300</sub>(1.5, 7.5) | $0.51 \pm 0.04$ | $0.50 \pm 0.03$ |

We note that both the pathway-reporter (nascent-mature or mature-protein) and dual reporter methods show slower convergence when copy numbers are low. Pathway reporters usually show fractionally slower convergence and fractionally more variation than a dual reporter, as suggested by the standard deviations in Tables 2 and 3. A more detailed exploration of convergence is given in the supplementary material.

### Measuring Noise from a Facultative (bursty) Gene

The most common mode of gene expression that is reported experimentally is burstiness [9,17,22,24,25,54], in which mRNA are produced in short bursts with periods of inactivity in between. One example is gene regulation via repression, which naturally leads to periods of gene inactivity. Here we consider a four-stage model of bursty gene expression, which incorporates both promoter switching and mRNA maturation; we denote this model by **M**_4_; see Fig. 4A (bottom left). This model has recently been considered in [37], where the marginal distributions are solved in some limiting cases. In this model, the gene switches probabilistically between an active state (A) and an inactive state (I), at rates *λ* (on-rate) and *µ* (off-rate), respectively. In the active state, nascent mRNA is synthesized according to a Poisson process at rate *K*_*N*_, while in the inactive state transcription does not occur. Nascent mRNA is spliced into mature mRNA at rate *K*_*M*_; this in turn is later translated into protein, with rate *K*_*P*_. Degradation of mRNA and protein occur independently of the promoter state at rates *δ*_*M*_ and *δ*_*P*_, respectively.

For this model we have three natural candidates for pathway reporters: (a) nascent and mature mRNA (b) mature mRNA and protein, and (c) nascent mRNA and protein reporters. Below we show that nascent mRNA–protein reporters yield consistently good estimates of the extrinsic noise contribution 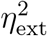 to the total gene expression noise, while nascent–mature and mature RNA–protein reporters are reliable in some restricted cases. We begin by showing that each of the reporter pairs (a), (b) and (c) satisfy the Noise Decomposition Principle. We then demonstrate computationally, that despite a lack of independence between these reporter pairs, the pathway-reporter method can still be used to decompose the total gene expression noise at the transcriptional level. Throughout, we let *X*_*N*_ denote the number of nascent mRNA, we let *X*_*M*_ denote the number of mature mRNA, and let *X*_*P*_ denote the number of proteins produced from the same gene. We also let **Z** = {*λ, µ, K*_*N*_, *K*_*M*_, *K*_*P*_, *δ*_*M*_, *δ*_*P*_}.

Folloqing [17] we assume that the transcription rate *K*_*N*_ is large relative to the other parameters. We further assume that the maturation rate *K*_*M*_ is large (i.e. *K*_*M*_ *> δ*_*M*_), which is supported by experiments [37]. Then, using the results of [17] and arguments similar to those in [37], it can be shown that the stationary averages for the nascent mRNA, mature mRNA and protein levels are given by,

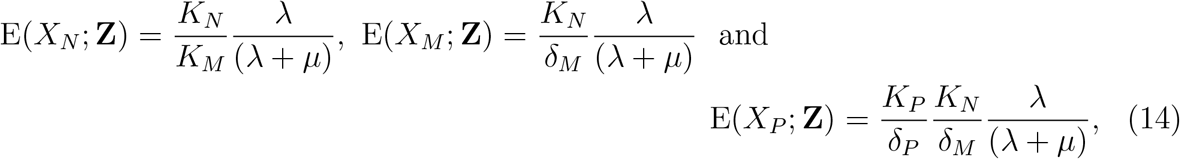

respectively. The averages for the mature mRNA and protein levels are given in [17] for the “three-stage model” of gene expression; see Fig. 4A (bottom right). The supplementary material provides more detail on how these expressions can be obtained.

We consider first the nascent-mature pathway reporters, case (a). From (14), it is easily seen that 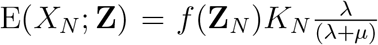 and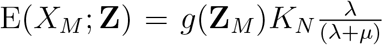, where 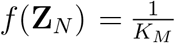 and 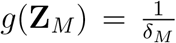. So the NDP holds, and the normalised covariance of E(*X*_*N*_; **Z**) and E(*X*_*M*_; **Z**) will identify the extrinsic noise on the transcription component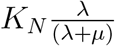. Consider now the mature-protein reporters, case (b). Again, we can see from (14) that E(*X*_*M*_; **Z**) = *f* (**Z**_*P*_) E(*X*; **Z**), where 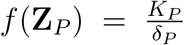. Thus the NDP holds, and so the normalised covariance of E(*X*_*M*_; **Z**) and E(*X*_*P*_; **Z**) will identify the total noise on E(*X*_*M*_; **Z**) (the extrinsic noise on *X*_*M*_). For the nascent-protein reporters, case (c), it is easy to see that 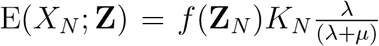, where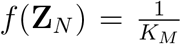, and 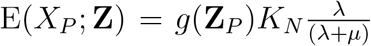, where 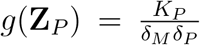. Thus, again the NDP holds, and the normalised covariance of E(*X*_*N*_; **Z**) and E(*X*_*P*_; **Z**) will identify the noise on the transcriptional component 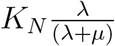.

In order for the pathway-reporter method to provide a close approximation to the extrinsic noise in cases (a), (b) and (c), we require that the normalised intrinsic contribution to the covariance is either zero or negligible. This condition will hold provided there is sufficiently small correlation between the reporter pairs. In the case of (prokaryotic) mRNA and protein, this lack of correlation has been been verified experimentally in E. Coli [55]. More generally, it is possible to provide an intuition about the conditions under which the lack of correlation might hold. The time series of copy numbers for each of nascent mRNA, mature mRNA and protein broadly follow each other, each with delay from its predecessor (Fig. 4B). Parameter values that reduce this delay will tend to increase correlation, and thereby increase the normalised intrinsic contribution to the covariance. The primary example of this effect is seen when *δ*_*p*_ approaches, or even exceeds *δ*_*m*_ (or for nascent-mature reporters, when *δ*_*M*_ approaches the maturation rate). A further contributor to high correlation between mRNA and protein, is when the system undergoes long timescale changes. In this situation the copy numbers tend to drop to very low values for extended periods. The primary parameter influencing this type of behaviour is the active-rate *λ*, specifically, when *λ* tends to 0 (Fig. 4C). An illustrative example of this can be seen by considering a Telegraph system in the limit of slow switching, which produces a copy number distribution that converges to a scaled Poisson-Bernoulli compound distribution: even without any extrinsic noise, the pathway reporter method will identify the *η*^2^ value of the corresponding scaled Bernoulli distribution. An extensive computational exploration of the parameter space (supplementary dataset S2) supports our intuition, though the strength of the effect varies across the three different reporter pairs. This is further corroborated by the heatmaps shown in Fig. 5, which for two fixed values of *µ* and a broad spectrum of *δ*_*p*_ and *λ* values, give the intrinsic term 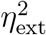 in (8), for fixed **Z**. Thus, the heatmaps provide an estimate for the overshoot error in the pathway-reporter approach. Note that blue pixels represent an overshoot estimate of less than 0.05. We obtain similar results for the intermediate case of *µ* = 10 (supplementary material, Fig. S4).

**Figure 5:**
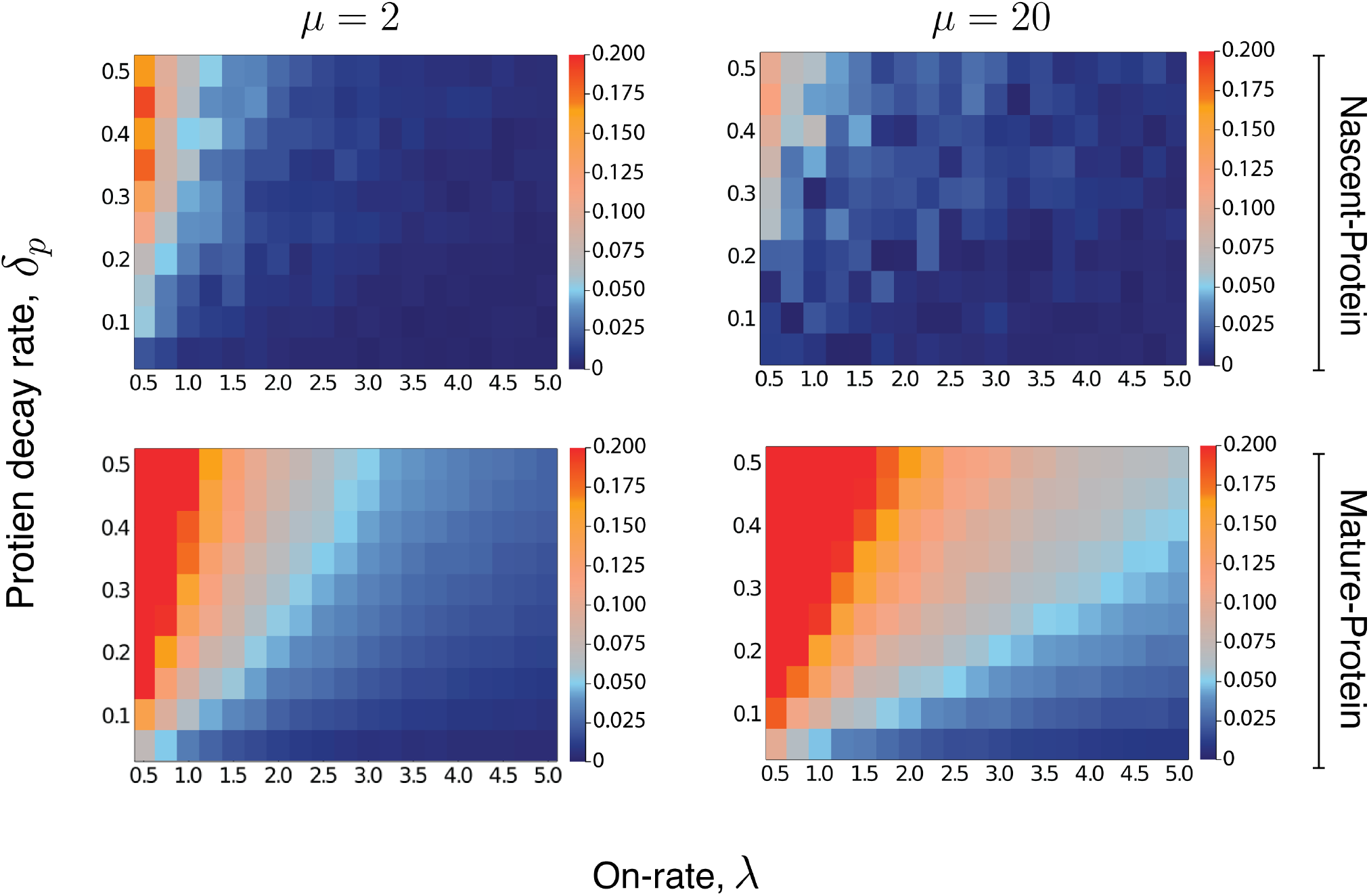
Heatmaps for the intrinsic contribution to the covariance, which estimates the level of overshoot in the pathway-reporter approach for the nascent-protein and mature-protein reporters; blue regions show an overshoot of less than *≃* 0.05. Here the intrinsic contribution is calculated using stochastic simulations of the model **M**_4_. In the left panels, the parameter *µ* = 2 is fixed, while *δ*_*p*_ and the on-rate *λ* are varied between 0.01 and 1 and 0.5 and 4, respectively. In the right panels, the parameter *µ* = 20 is fixed, while *δ*_*p*_ and the on-rate *λ* vary between 0.01 and 1, and 1 and 10, respectively. In all cases, the parameters are scaled so that *δ*_*M*_ = 1. The maturation rate is fixed at 20, with the parameters *K*_*N*_ and *K*_*P*_ are chosen to produce a mean protein level of 1000, a mean nascent mRNA level of 5 and a mean mature mRNA level of 50.

For nascent-protein reporters, the normalised intrinsic contribution to the covariance is satisfactorily small (less than 0.05) for (a) high values of *δ*_*p*_ in unison with (b) low values of *λ* (less than 1, though lower values are acceptable if *δ*_*p*_ is small). The assumption (a) that *δ*_*P*_ *< δ*_*M*_ is known to be true for a large number of genes, and is justified by the difference in the mRNA and protein lifetimes. While there is of course variation across genes and organisms, values of *δ*_*P*_ *≤* 0.5*δ*_*M*_ and even *δ*_*P*_ *≤* 0.2*δ*_*M*_ seem reasonable for the majority of genes. In E. Coli [55] and yeast [56], for example, mRNA are typically degraded within a few minutes, while most proteins have lifetimes at the level of 10s of minutes to hours. For mammalian genes [53], it has been reported that the median mRNA decay rate *δ*_*M*_ is (approximately) five times larger than the median protein decay rate *δ*_*P*_, determined from 4, 200 genes. Assumption (b) requires that the gene is sufficiently active. In a recent paper by Larsson et al. [25], the promoter-switching rates *λ, µ*, and the transcription rate *K* of the Telegraph model are estimated from single-cell data for over 7, 000 genes in mouse and human fibroblasts. Of those genes with mean mRNA levels greater than 5, we found that over 90% have a value for *λ* of at least 0.5, and over 65% have a value for *λ* greater than 1. In Cao and Grima [37], a comprehensive list of genes (ranging from yeast cells through to human cells) with experimentally-inferred parameter values are sourced from across the literature (see Table 1 in [37]). After scaling the parameter values of the 26 genes reported there, we find that around 88% have a value for *λ* of at least 0.5, and approximately 58% have a value for *λ* greater than 1. Thus, the heatmaps given in Fig. 5 (top panel) suggest that nascent-protein reporters will provide a satisfactory estimate of the extrinsic noise level for a substantial fraction of genes.

The mature nRNA–protein reporters are less reliable, with the requirement of slow protein decay and higher on-rate being more pronounced than for the nascent mRNA– protein reporters; this is evident from Fig. 5 (second panel). The performance of the nascent–mature reporter is of course independent of *δ*_*p*_, but is only viable in the case of a large on-rate (see Fig. S5 of the supplementary material).

We tested our approach for each of the pathway reporter pairs (a), (b) and (c) against dual reporters using stochastic simulations. Table 4 shows the results from six simulations across a spectrum of behaviours from moderately slow switching, to fast switching as well as a range of levels of burstiness. For each of the parameters *µ, λ, K*_*P*_, *δ*_*P*_ we selected a scaled Beta(5, 6) distribution, with squared coefficient of variation *η*^2^ = 0.1; the scaling is chosen in each case to achieve a mean value equal to the parameter value given in Table 4. It is routinely verified that scaling these distributions does not change the value of *η*^2^. The parameter *K*_*N*_ is given the noise distribution Beta(3, 6), which has a slightly higher coefficient of variation *η*^2^ = 0.2. In order to achieve direct benchmarking against the dual reporters, the parameter *K*_*M*_ is fixed at 10. This is because the nascent-protein pathway reporter estimates noise on the value of 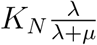, while the mature mRNA dual-reporter measures noise on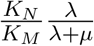, and these coincide only when *K*_*M*_ is fixed. The mean values of *K*_*N*_ and *K*_*P*_ are chosen to achieve approximate average nascent mRNA levels, mature mRNA levels and protein levels at 5, 50 and 1000 respectively, given the chosen values of *λ, µ, δ*_*P*_.

**Table 4:** A comparison of the pathway-reporter method and dual-reporter method for bursty expression. Here PR (NP) gives the results of the nascent and protein pathway reporters, PR (MP) gives the results of the mRNA and protein reporters, while DR (Mat) gives the results of the dual reporters calculated from the mature mRNA. We consider noise on all of the parameters except for *δ*_*M*_ and *K*_*M*_; see discussion in main text. The values given are the average of 100 simulations, each calculated from 500 copy number samples, and the errors are one standard deviation. Our theory predicts that pathway-reporters will identify the noise at both the promoter level (*λ, µ*) and transcriptional level (*K*_*N*_, *δ*_*m*_); the total extrinsic noise in each case is given by 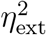. As before, the noise distribution parameters are chosen to produce an average nascent mRNA copy number of 5 and an average mature mRNA copy number of 50, and an average number of 1000 proteins.

| Mean |  |  |  |  | Simulation |  |  |
| --- | --- | --- | --- | --- | --- | --- | --- |
| $\lambda$ | $\mu$ | $K_N$ | $K_P$ | $\delta_P$ | PR (MP) | PR (NP) | DR (Mat) |
| 0.5 | 1 | 150 | 2 | 0.1 | $0.46 \pm 0.06$ | $0.38 \pm 0.07$ | $0.32 \pm 0.07$ |
| 1 | 2 | 150 | 2 | 0.1 | $0.39 \pm 0.05$ | $0.34 \pm 0.07$ | $0.32 \pm 0.05$ |
| 1 | 20 | 1050 | 2 | 0.1 | $0.66 \pm 0.15$ | $0.52 \pm 0.22$ | $0.50 \pm 0.15$ |
| 2 | 2 | 100 | 6 | 0.3 | $0.35 \pm 0.04$ | $0.29 \pm 0.05$ | $0.27 \pm 0.03$ |
| 2 | 20 | 550 | 6 | 0.3 | $0.61 \pm 0.09$ | $0.47 \pm 0.15$ | $0.47 \pm 0.09$ |
| 10 | 10 | 100 | 6 | 0.3 | $0.29 \pm 0.03$ | $0.27 \pm 0.04$ | $0.27 \pm 0.02$ |

The results for the nascent mRNA–protein reporters, case (c), given in Table 4 show comparable performance to dual reporters, with only modest overshoot; even in the worst performing case of *λ* = 0.5, *µ* = 1 the result of the pathway reporters is within one standard deviation, in a very tight distribution. The error heatmaps of Fig. 5 provide an accurate estimate of the overshoot in the nascent-protein results in Table 4. As an example, the first row is most closely matched by the heatmap at top left of Fig. 5, which at *λ* = 0.5 and *δ*_*P*_ = 0.1 is suggestive of an error around the boundary between blue and red (around 0.06). The same accuracy is obtained for the other rows. As predicted, the mature-protein reporters show significantly more overshoot, especially with the less active genes. Improved accuracy can again be obtained by subtracting the estimated overshoot given in the error heatmaps from the obtained value. Thus for example, the error heatmap for *µ* = 2 (Fig. 5 lower left) gives an error approximately 0.07 for *λ* = 1, *δ*_*P*_ = 0.1, which agrees very closely to the actual overshoot of 0.07 shown in the corresponding row of Table 4. An overshoot of approximately 0.06 is suggested by the heatmap for *µ* = 2, when *λ* = 2, *δ*_*P*_ = 0.3, which leads to a correction from 0.35 in Table 4 to a value of 0.29. This is quite consistent with the dual reporter benchmark of 0.27. As expected (based on Fig. S5 in the supplementary material), nascent-mature reporters do not perform well on bursty systems except for high *λ* and so the values are not included in Table 4; only in the case of *λ* = *µ* = 10 does the result begin to approach the dual reporter value, returning 0.32 *±* 0.03 (Fig. S5 in the supplementary material).

## Discussion

Despite the proliferation of experimental methods for single-cell profiling, the ability to extract transcriptional dynamics from measured distributions of mRNA copy numbers is limited. In particular, the multiple factors that contribute to mRNA heterogeneity can confound the measured distribution, which hinders analysis. Theoretical innovations that allow us to quantify and help identifying the causes of these observable effects are therefore of great importance. In this work, we have demonstrated, through a series of mathematical results, that it is impossible to delineate the relative sources of heterogeneity from the measured transcript abundance distribution alone: multiple possible dynamics can give rise to the same distribution. Our approach involves establishing integral representations for distributions that are commonly encountered in single-cell data analysis, such as the negative binomial distribution and the stationary probability distribution of the Telegraph model. We show that a number of well-known representations can be obtained from our results. A particular feature of our non-identifiability results is that population heterogeneity inflates the apparent burstiness of the system. It is therefore necessary to collect further information, beyond measurements of the transcripts alone, in order to constrain the number of possible theoretical models of gene activity that could represent the system. In particular, additional work may be required to determine the true level of burstiness of the underlying system.

We have developed a theoretical framework for estimating levels of extrinsic noise, which can assist in resolving the non-identifiability problems. The dual reporter method of Swain et al. [4] already provides one such approach; but it is experimentally challenging to set up in many systems, and requires strictly identical and independent pairs of gene reporters. Our Noise Decomposition Principle directly generalises the theoretical underpinnings of the dual reporter method; here we have used it to identify a practical approach —the pathway-reporter method— for obtaining the same noise decomposition. Our approach allows us to use measurements of two different species from the transcriptional pathway of a single gene copy instead of having to set up a more cumbersome dual reporter assay. The accuracy of the pathway-reporter method is provably identical for constitutive gene expression, and in the case of nascent-mature mRNA reporters, the measurements are readily obtainable from current single-cell data [13, 42, 57]. For bursty systems, the method in general provides only an approximation of the extrinsic noise. We are, however, able to demonstrate computationally, that one of the proposed pathway reporters provides a satisfactory estimate of the extrinsic noise for most genes. The other pathway reporters also provide viable estimates of the extrinsic noise in some cases.

Despite the generality of our theoretical contribution, our pathway-reporter approach has some caveats. In particular, the approach relies on the assumption that extrinsic noise sources act independently. Experimentally, however, these may be correlated. For example, it has been suggested [58, 59] that the transcription and translation rates in E. coli anticorrelate. Additional work is required to determine degree to which the independence of noise sources is a reasonable assumption.

Recent developments in single-cell profiling now allow simultaneous measurements of transcripts and proteins in thousands of single cells [60–62]. As discussed in [14], experimental improvements that would additionally allow measurements of nascent transcripts, coupled with theoretical developments such as those presented here, will allow for identification of noise sources on a genome-wide scale. Our work reveals that extrinsic noise distorts the multivariate copy number distribution of the different species in the gene expression pathway. We have exploited this to yield reliable estimates of noise strength, which we are confident will assist in setting better practices for model fitting and inference in the analysis of single-cell data. A more nuanced analysis of this multivariate distribution may offer even further insight into model and noise identifiability.

## Acknowledgments

We gratefully acknowledge Rowan D. Brackston helpful discussions in the early stages of this research. We also wish to thank Arjun Raj for providing valuable feedback on this work. L.H. and M.P.H.S. were supported by the University of Melbourne DVCR fund.

## Data Availability

Simulations of the models used in the paper are performed using Gillespie’s Stochastic Simulation Algorithm (SSA) implemented in Julia. The simulation code is available in the GitHub repository https://github.com/leham/PathwayReporters. The data used in the paper are provided in the supplementary datasets.

## Author Contributions

L.H. and M.J. conceptualised the research, with support from M.P.H.S. The theoretical framework was developed by L.H. Formal analysis was conducted by L.H. and M.J. Original draft was written by L.H., with editing and review contributed by M.J. and M.P.H.S. All authors provided critical feedback and helped shape the research.

## Supplementary Text

### Derivations of the Non-Identifiability Results

In the main text we provide a number of examples of when the compound distribution does not have a unique representation. We here provide the full details of the derivations of these results.

#### Bursty Expression: Telegraph Representation

Consider first a Telegraph distribution 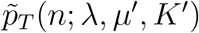 and the probability density function of the scaled beta distribution Beta_*K′*_(*λ* + *µ, µ*^*′*^ *− µ*), given by

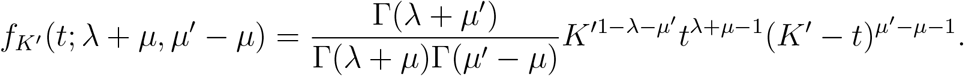

Note that this distribution has support [0, *K*^*′*^]. In the main text we claim that the Telegraph distribution 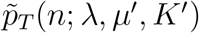 can be obtained from compounding a Telegraph distribution by a scaled beta distribution with pdf *f* (*t*; *λ* + *µ, µ*^*′*^ *− µ*). In other words, that 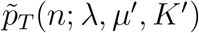 can be written as:

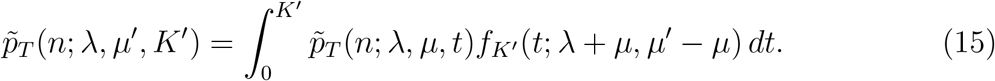

This is Eq. (3) in the main text. Starting from the right hand side of (15), we have

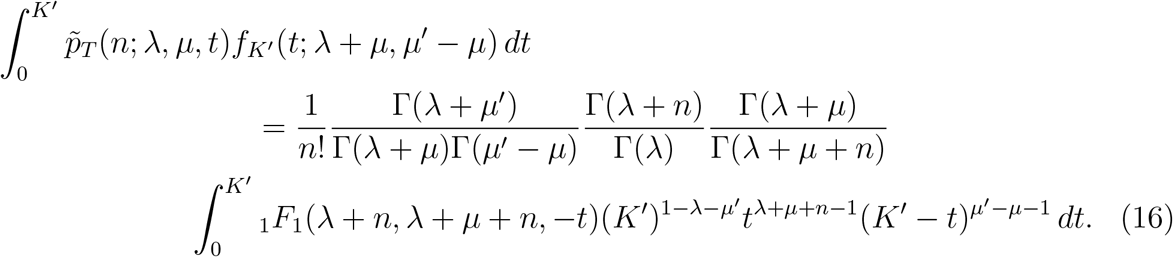

Substituting *t* = *K*^*′*^*T* and simplifying, the right hand side of (16) becomes

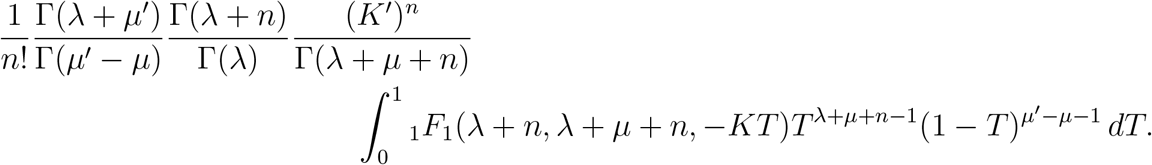

Now, using the integral representation [63][13.4.2] of the confluent hypergeometric function (with *a* = *λ* + *n, b* = *λ* + *µ*^*′*^ + 1 and *c* = *λ* + *µ* + 1), this becomes

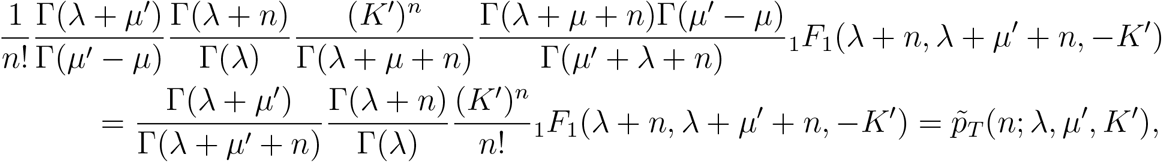

which is the left hand side of (15). Hence, we have that (15) holds.

#### Instantaneously Bursty Expression: Negative Binomial Representations

We consider first the representation given in Eq. (4) in the main t ext. Recall that we let 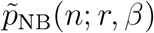 denote the probability mass function of a 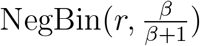 distribution, where *β >* 0. In the main text we claim that for any negative binomial distribution of the form NegBin 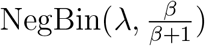 (where *β >* 0) we have,

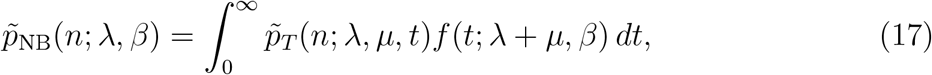

where *f* (*t*; *λ* + *µ, β*) is the probability mass function of a Gamma(*λ* + *µ, β*) distribution. Beginning with the right hand side of (17) we have,

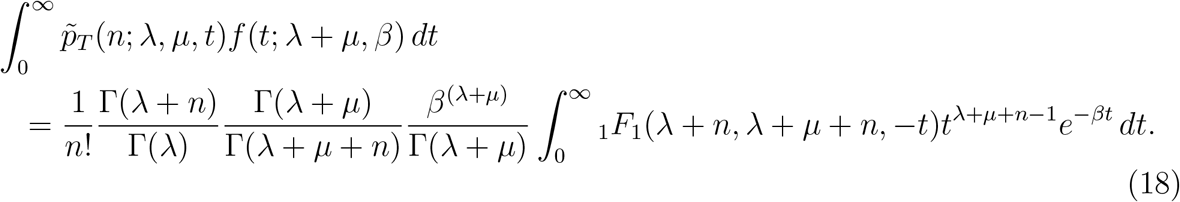

We now apply the following identity given in [64][Appendix 1] with *a* = *λ*+*n, b* = *λ*+*µ*+*n, k* = *−*1, *d* = *λ* + *µ* + *n* and *h* = *β*.

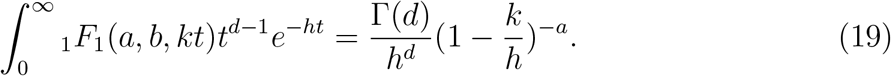

So the left hand side of (18) becomes

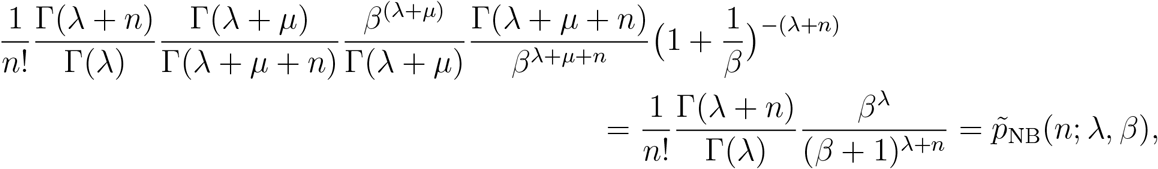

which is the right hand side of (17). Hence, we have that (17) holds. We now consider the representation given in Eq. (5) of the main text. Here we claim that any negative binomial distribution of the form 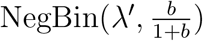 (where *b >* 0) can be written as,

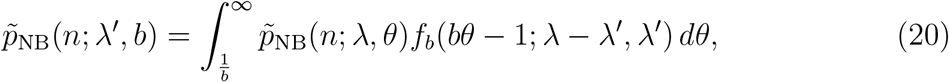

where *f*_*b*_(*bθ−*1; *λ−λ*^*′*^, *λ*^*′*^) is the probability mass function of a scaled beta prime BetaPrime_*b*_(*λ − λ*^*′*^, *λ*) distribution, where *b >* 0 and *λ > λ*^*′*^. Starting from the right hand side of (20), we have

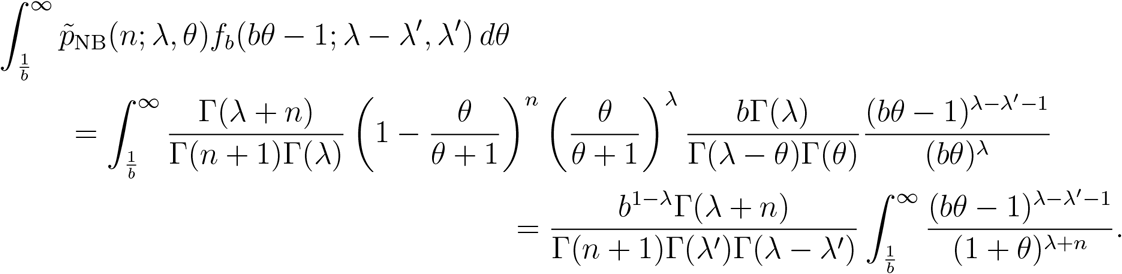

Now substituting *H* = *bθ −* 1 and simplifying, we obtain

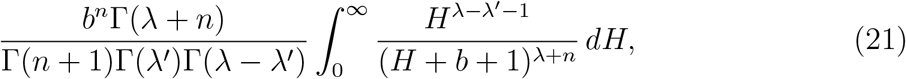

and letting *y* = *b* + 1 and simplifying we have that

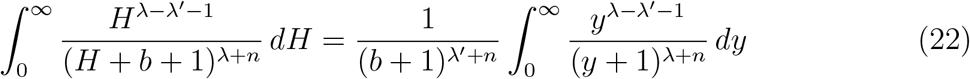

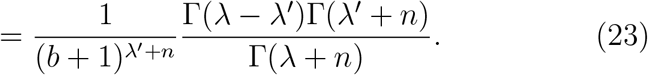

Here we used the fact that the integral in the variable *y* is the probability density function of a BetaPrime(*λ − λ*^*′*^, *λ*^*′*^ + *n*) distribution. Thus, Eq. 21 simplifies to

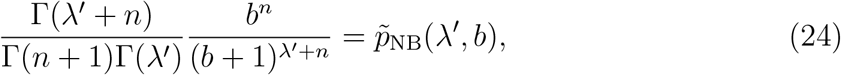

which is the right hand side of (20). Thus, we have that (20) holds.

### Proof of the Noise Decomposition Principle

Here we provide the proof of the Noise Decomposition Principle given in the main text. For convenience we restate the principle below.

#### The Noise Decomposition Principle (NDP)

Assume that there are measurable functions *f, g* and *h* such that E(*X*; **Z**) and E(*Y*; **Z**) split across common variables by way of E(*X*; **Z**) = *f* (**Z**_*X*_)*h*(**Z**^*′*^) and E(*Y*; **Z**) = *g*(**Z**_*Y*_)*h*(**Z**^*′*^). Then provided that the variables *Z*_1_, …, *Z*_*m*_ are mutually independent, the normalised covariance of E(*X*; **Z**) and E(*Y*; **Z**) will identify the total noise on *h*(**Z**^*′*^) (i.e.,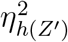).

Consider first the covariance of E(*X*; **Z**) and E(*Y*; **Z**). We have

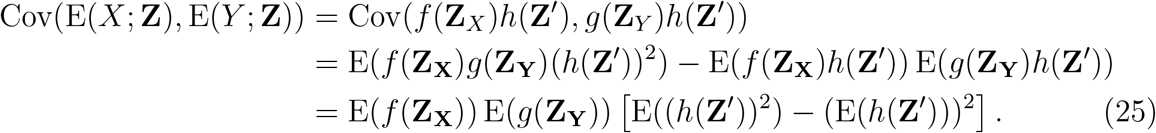

Note that here we used the fact that the variables in **Z** = (*Z*_1_, …, *Z*_*n*_) are mutually independent. Now using the fact that E(*X*) = E(E(*X*; **Z**)) and E(*Y*) = E(E(*Y*; **Z**)) (the Law of Total Expectation), and then normalising we obtain

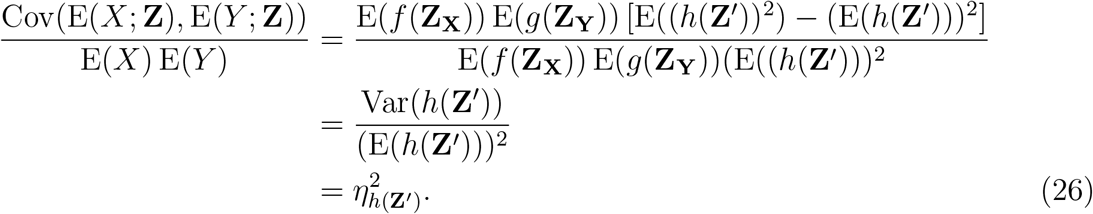

Hence, under certain conditions the normalised covariance of E(*X*; **Z**) and E(*Y*; **Z**) will identify the total noise on *h*(**Z**^*′*^).

## Pathway Reporters

### Justification for Simulating Only One Copy of the Gene

Our simulations and theory have been based over reporters from a single gene copy, whereas in practice there may be multiple copies of the gene that contribute to the overall mRNA. If there are mechanisms in place to distinguish the mRNA or protein of one gene copy from another, then the theory and analysis we have developed in main paper holds without change. In the case where it is not possible to distinguish mRNA and protein (respectively) from the multiple gene copies, then we now observe that the general theory continues to hold, provided there is independence between the gene copies; this assumption has been verified experimentally [13]. Here we demonstrate this in the case of two gene copies, though the general case for more than two genes is essentially identical but is notationally cumbersome. We are considering a situation where the variable *X* in the Noise Decomposition Principle is the sum of two independent variables *X*_1_, *X*_2_ and *Y* is the sum of two independent variables *Y*_1_, *Y*_2_. We assume common dependence on the environmental variables **Z** so that *E*(*X*_1_; **Z**) = *E*(*X*_2_; **Z**), *E*(*Y*_1_; **Z**) = *E*(*Y*_2_; **Z**). Using these equalities and the independence of *X*_1_, *X*_2_ and *Y*_1_, *Y*_2_ in *X* = *X*_1_ + *X*_2_, *Y* = *Y*_1_ + *Y*_2_, we find the numerator of

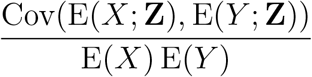

is simply 4 Cov(E(*X*_1_; **Z**), E(*Y*_1_; **Z**)), while the denominator is 4 E(*X*_1_) E(*Y*_1_). Thus the noise decomposition coincides with that for the single copy *X*_1_, *Y*_1_. Further work may be required to consider systems where there is independence between, or there is significant deviations in the gene copies.

### Upper Bound for the Intrinsic Contribution to the Covariance: Constitutive Expression

In the main text, we claim that the error in the pathway-reporter approach in the case of mRNA-protein reporters is negligible (i.e. the error is *≪* 1). Our argument relies on the expression given in Eq. 13 in the main text. We here provide full details of the derivation of this expression. First let *X*_*m*_ and *X*_*p*_ be the number of mRNA and protein produced from the same constitutive gene modelled by the “two-stage” model, **M**_2_ (see Figure 3A (top right) of the main text). Also, let **Z** = {*K*_*m*_, *δ*_*m*_, *K*_*p*_, *δ*_*p*_}.

We restate here the expression for the intrinsic contribution to the covariance of *X*_*m*_and *X*_*p*_ given in Eq. 13 of the main text.

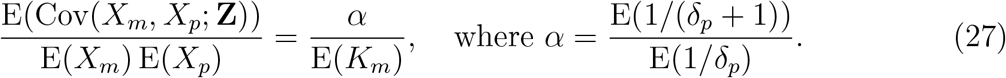

We require the following expressions for the stationary mean mRNA level and protein level of the two-stage model [46, 48].

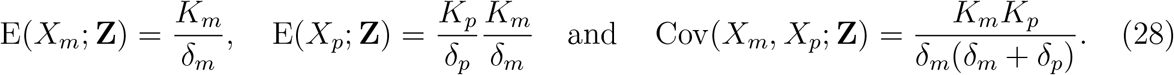

Assuming that *δ*_*m*_ is fixed across the cell-population, and all parameters are scaled so that *δ*_*m*_ = 1, it follows from (28), that

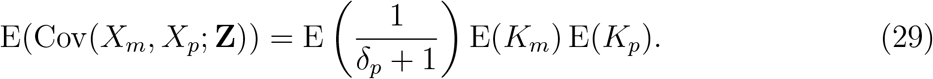

Using the Total Law of Expectation, we also have,

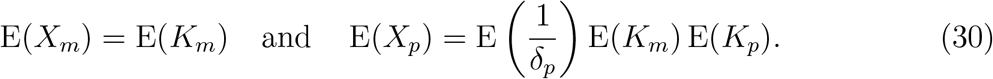

Thus, it follows that

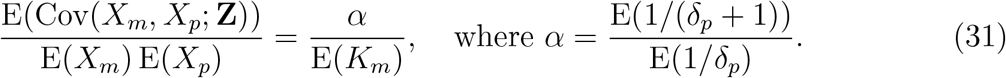

### Determination of the Marginal Means in the Model M4

In order to establish the Noise Decomposition principle in the case of bursty gene expression, we rely on expressions for the marginal stationary means of the model **M**_4_; see Fig. 3A (bottom left) and the associated caption for more details of this model. To derive the marginal means for nascent and mature mRNA and for protein (Eq. 14 of the main text), we first observe that the nascent mRNA population may treated identically to that of mRNA in general (that is, no distinction between nascent and mature), as in [26], except that mRNA decay is replaced by the sum of decay and maturation. As in the work of Cao and Grima [37], the assumption of fast maturation allows us to ignore decay completely in the nascent phase, so that the marginal distribution is identical to that of [26], except with decay replaced by maturation. This leads to a marginal nascent mRNA mean of

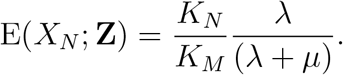

The marginal means for mature mRNA and protein are derived in [17] under the assumption that the transcription rate parameter *K*_*N*_ is large relative to the other parameters. The expressions are given by

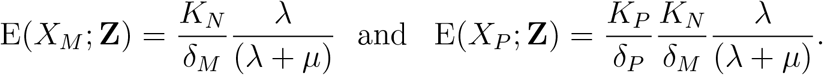

Formally, the marginal means in [17] are for the three-stage model **M**_3_, which ignores the downstream processing of mRNA, such as splicing. The assumption of fast maturation however, justifies the treatment of the nascent phase of mRNA as an ephemeral step within the Poissonian modelling of mRNA transcription.

There are a number of possibly compounding assumptions on the parameters here, but simulations show that there is a lot of tolerance, with even only moderate maturation and transcription still returning sample means consistent with the formulas.

### Measuring Noise from an Instantaneously Bursty Gene

Here we use the simple stochastic model of [65] that includes both instantaneous transcriptional bursting and mRNA maturation. This model can be obtained as a limiting case of the model **M**_4_ (Fig. 3A (top left) in the main text), where the off-rate *µ* has tended toward infinity, while the on-rate *λ* remains fixed. Under this condition, transcription is rare enough to be considered instantaneous, leading to “bursts” of transcriptional activity. The intensity of the bursts *M* is known to be geometrically distributed with mean burst size 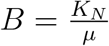.

If we let *X*_*N*_ and *X*_*M*_ denote the number of nascent and mature mRNA, respectively, then according to [65], the steady-state marginal means and covariance are given by

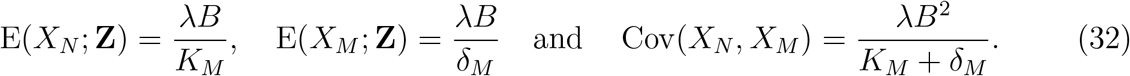

It is easily seen from (32) that E(*X*_*N*_; **Z**) = *f* (**Z**_*N*_)*Bλ* and E(*X*_*M*_; **Z**) = *g*(**Z**_*M*_)*Bλ*, where 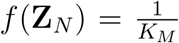 and 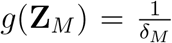. Thus, the Noise Decomposition Principle holds. Using arguments similar to those given in Section above, it is straightforward to derive the following expression for the error in the pathway reporter approach:

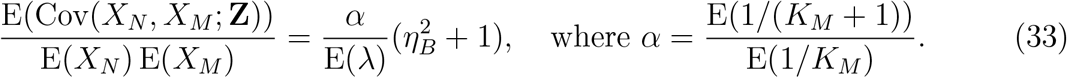

We can expect that *α ≈* 1, so that unless the burst frequency is substantial, that is *λ >>* 1, the overshoot in the the nascent-mature pathway-reporter approach is significant.

Convergence for the mature-protein reporters is considered in Figure S2. Here Eq. (13) shows that we should expect an overshoot in comparison to the mature-mature dual reporter. We have again compared a low output gene with a high output gene, and with all parameters (except *δ*_*M*_) experiencing noise. The transcription is again given scaled Beta(3, 6) distribution, having squared coefficient of variation 0.2, with scaling to achieve *E*(*K*_*N*_) = 5 in low output and *E*(*K*_*N*_) = 50 in the high output. The noise distribution for *δ*_*P*_ is a Beta(8, 6) distribution, scaled to achieve a mean value of 0.2. Computational sampling from 10^7^ samples finds the value of 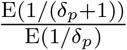 as approximately 0.1573. The high output gene then expects an overshoot of 0.003146 from the mature-protein reporters, while the low output gene expects an overshoot of approximately 0.03146. Figure S2.A shows comparison of convergence over the first 2000 samples, with the theoretical values accommodating the calculated overshoot. Figure S2.B shows the same over 20, 000 samples; this time two low-output two high-output genes are considered, so that the variation around the expected long term value can be seen. It is evident that reasonable estimates are given after a relatively modest number of samples, but there is a very long delay to a highly accurate convergence for the pathway reporters in the low output gene.

### Convergence of Pathway and Dual Reporters

Convergence of both pathway reporters and dual reporters in low output genes is slower than for high output genes, and this affect is most pronounced for reporters taken from nascent mRNA. Figure S1 compares convergence in some low output genes and high output genes for nascent-mature pathway reporters and the mature-mature dual reporter, in the case of constitutive expression. From Eq. (11) and the NDP, these should measure precisely the extrinsic noise on transcription. In the low output gene, both the dual reporter and pathway reporters are yet to show accurate measurement of the overall extrinsic noise after 1000 samples (Figure S1.A), though they are providing rough estimates after as little as a few hundred samples. Pathway reporters for nascent-mature typically exhibit slightly slower convergence than mature-mature dual reporter; the difference is marginal, but can be seen for both the high output gene (compare the values after 500 samples) and the low output gene (compare the values after 1000 samples). In this figure, all parameters (except the reference parameter *δ*_*M*_) were given scaled Beta distribution noise. The noise on *K*_*N*_ is a Beta(3, 6) distribution scaled to achieve *E*(*K*_*N*_) = 5 in the low output case and *E*(*K*_*N*_) = 50 in the high output case. The squared coefficient of variation is 0.2, and is shown in green in the figure.

**Figure S1:**
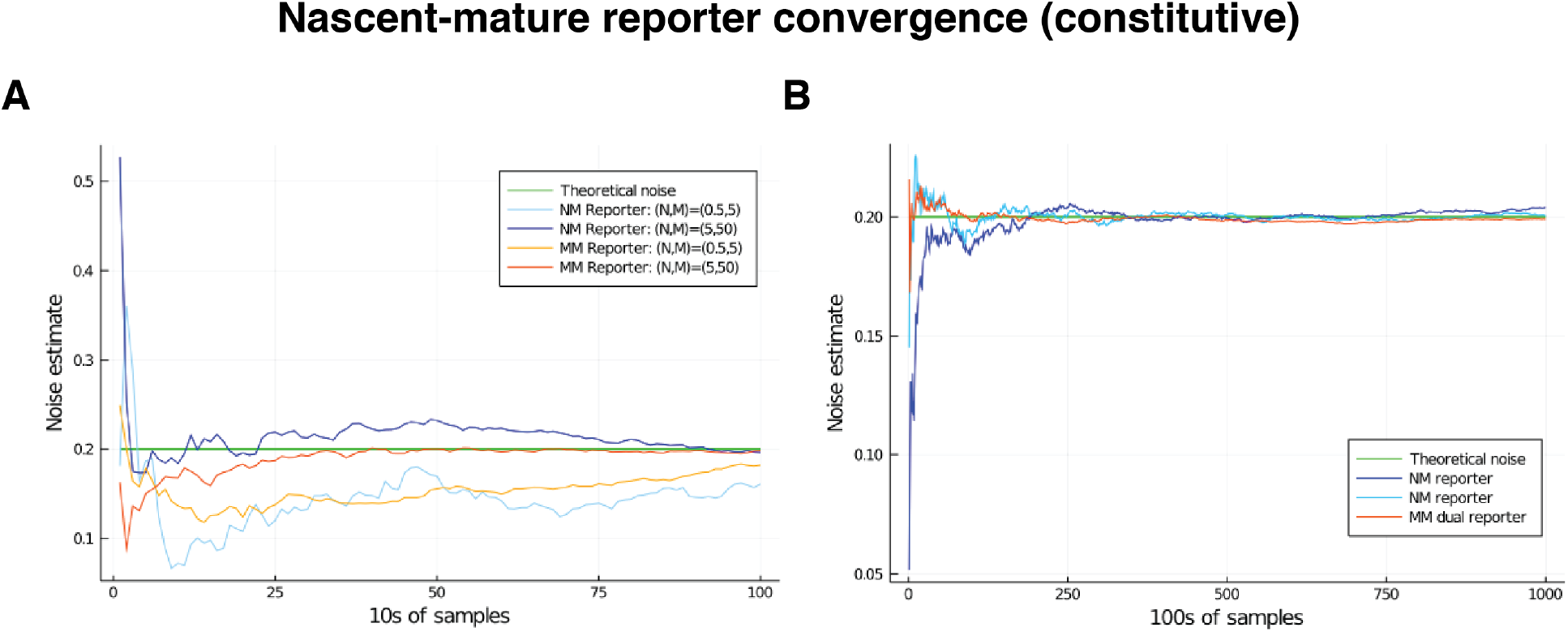
Comparison of convergence of *η*^2^ estimates for low and high mRNA levels by way of nascent-mature reporters and mature-mature reporters. Low level corresponds to mean nascent mRNA level of 0.5, and mean mature mRNA level of 5. High level corresponds to mean nascent mRNA level of 5, and mean mature mRNA level of 50. In both cases the simulated genes are constitutive and noise is on all parameters except for *δ*_*M*_ = 1. The green line gives the squared coefficient of variation for *K*_*N*_, set to 0.2, which is the value the various reporters are expected to estimate. A. Convergence of the *η*^2^ estimate over the first 2000 samples in the low- and high-output genes. B. Convergence of the *η*^2^ estimate over 100,000 samples in the low-output gene only.

**Figure S2:**
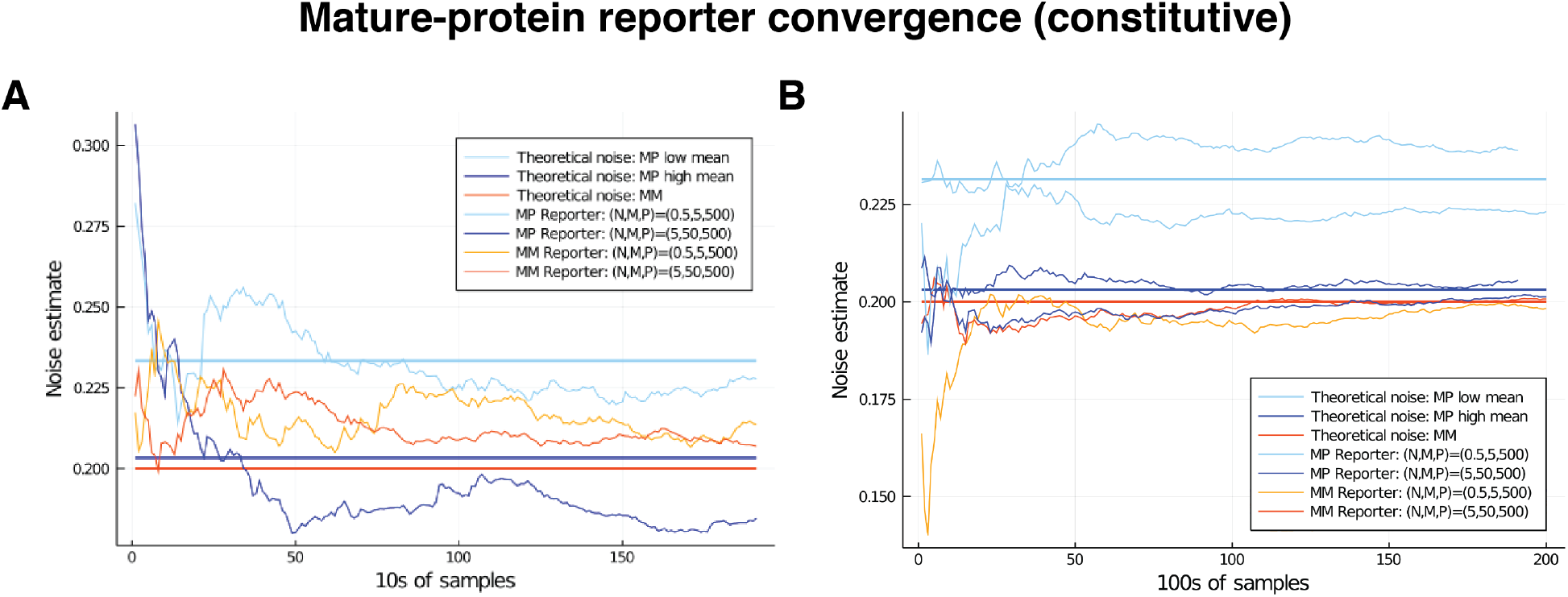
Comparison of convergence for low and high mRNA levels by way of mature-protein and mature-mature reporters. Low level corresponds to mean nascent mRNA level of 0.5, and mean mature mRNA level of 5. High level corresponds to mean nascent mRNA level of 5, and mean mature mRNA level of 50. In both cases the simulated genes are constitutive and noise is on all parameters except for *δ*_*M*_ = 1. The noise on *K*_*N*_ has squared coefficient of variation equal to 0.2, which is shown as the red horizontal line. Our theory shows that mature-protein reporters will return an overshoot that is negligible in the high-output gene (the blue horizontal line), but larger in the low output gene (light blue horizontal line); these values are calculated in the text. A. Comparison of convergence for low and high mRNA levels over the first 2000 samples. B. Convergence of the *η*^2^ estimate over 20,000 samples in the case of the low-output gene only. Two examples of each are given, to show the variation in behaviour.

**Figure S3:**
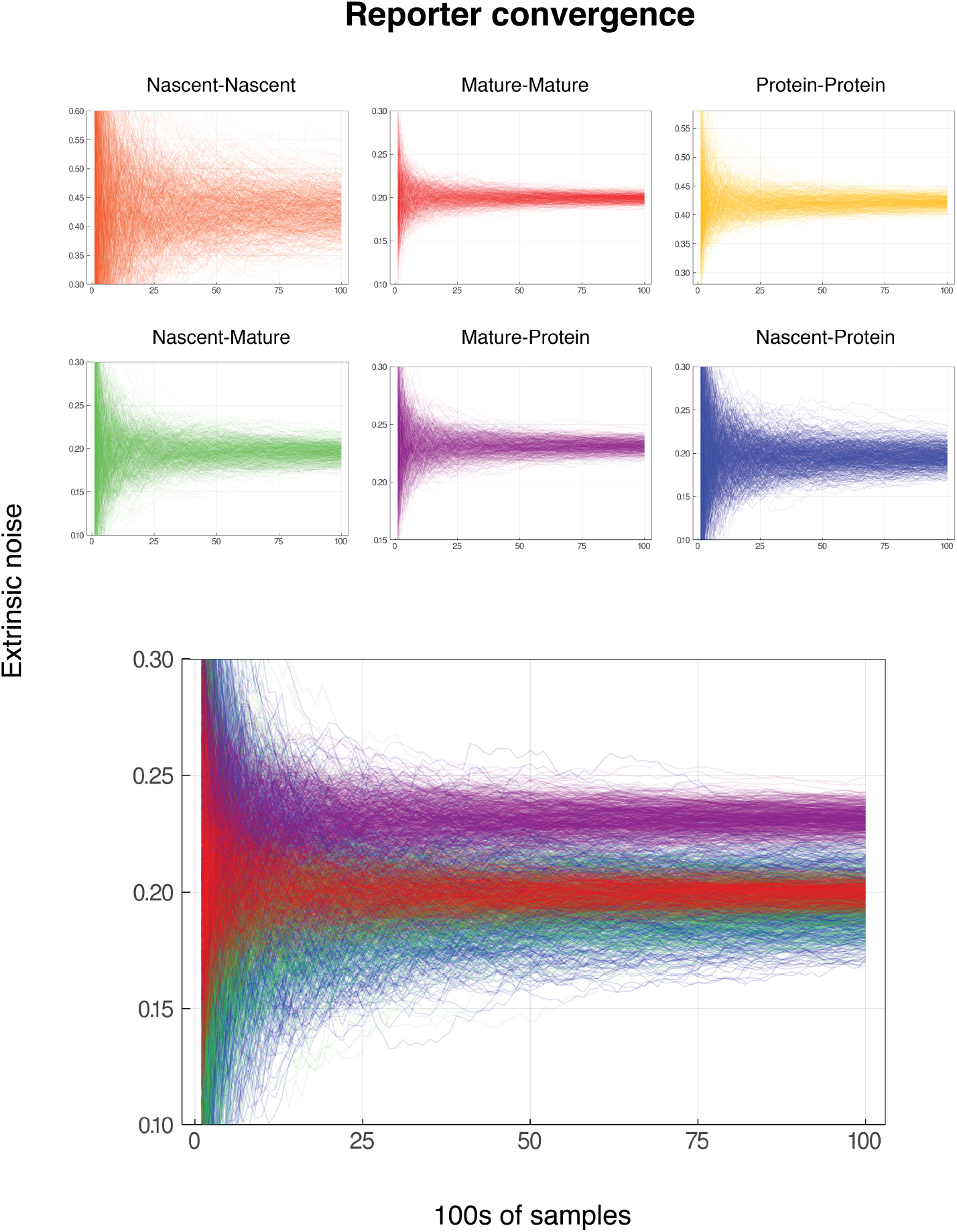
Convergence for reporter pairs for low gene activity. In each case, the mean nascent mRNA level is 0.5, the mean mature mRNA level of 5, and the mean protein level is 500. The simulated genes are constitutive and noise is on all parameters except for *δ*_*M*_ = 1. Each graph shows the convergence of 600 individual reporter simulations, for each combiantion of reporters from nascent mRNA, mature mRNA and protein. Each reporter simulation is from 10,000 samples, with the reporter estimates calculated at intervals of 100. The noise on *K*_*N*_ has squared coefficient of variation equal to 0.2, which should be identified by both the nascent-mature reporter and mature-mature dual reporter. As in Figure S2, the mature-protein reporter should converge to an estimate of approximately 0.2315. Nascent-nascent and protein-protein reporters identify combined noise on more parameters, so do not converge to 0.2. The lower graph shows each of nascent-mature, mature-mature, mature-pro4t0ein and nascent-protein in the same plot for direct comparison.

**Figure S4:**
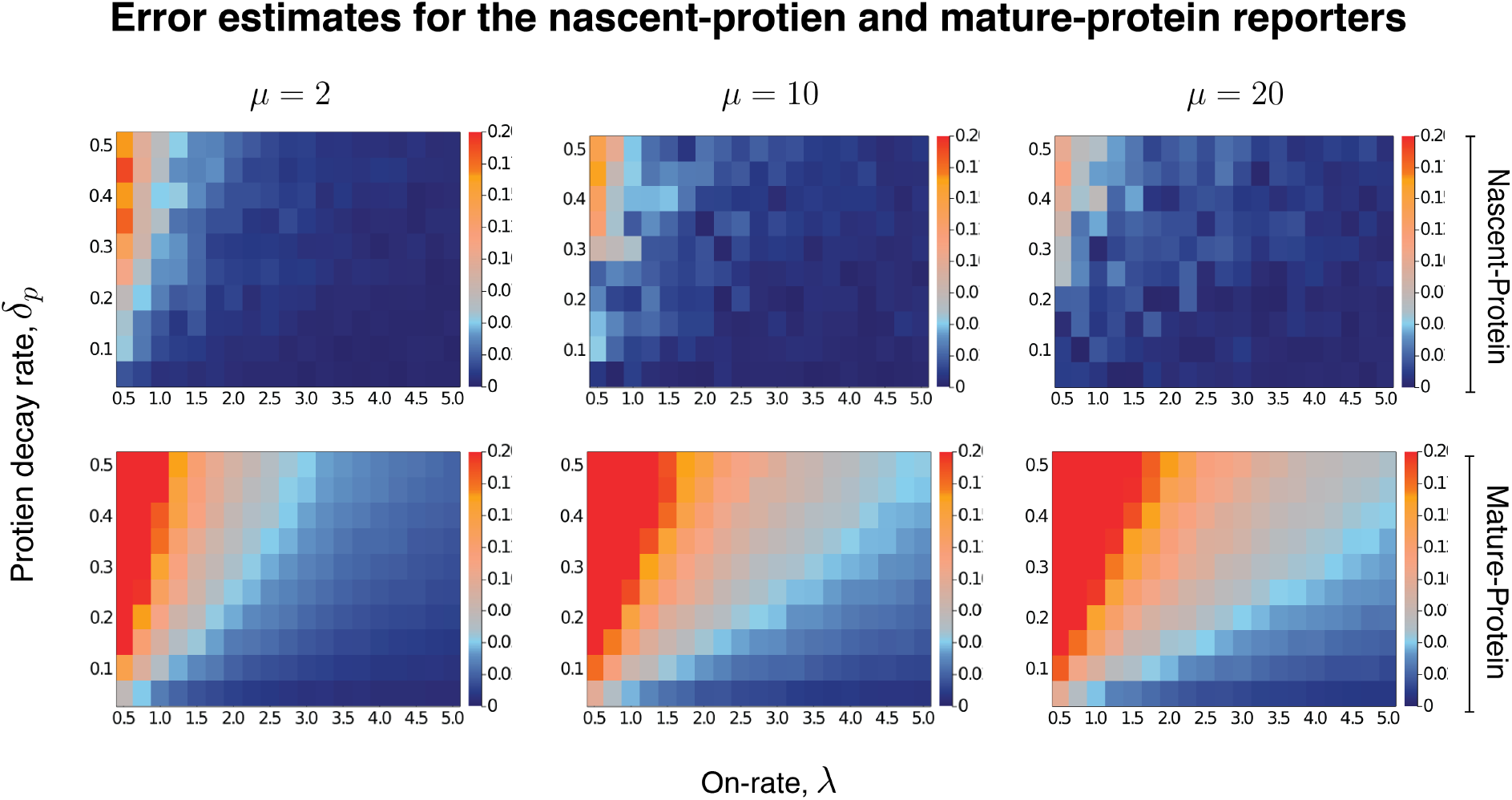
An extension of Fig. 3D of the main text. The heatmaps shown estimate the level of overshoot in the pathway-reporter approach, for both the nascent-protein and mature-protein reporters. For more details refer to the caption given in Fig. 3 in the main text. Here we include the additional case of *µ* = 10. The remaining parameter values are the same as in the main text. Each individual pixel is generated from a sample of size 3000, though there is still some instability in the convergence for the nascent-protein reporter, particularly as the overshoot estimation starts to increase, and particularly as *µ* is larger. To produce more accurate values, the case of *µ* = 2 was averaged over two full experiments while *µ* = 20 was averaged over three. This was also done for the mature-protein reporter, however for these images there was almost no visible difference between the various runs of the experiment and their averages. Each of the three *µ* values takes roughly 7–10 hours of computation, depending on lead in time before sampling within a simulation.

**Figure S5:**
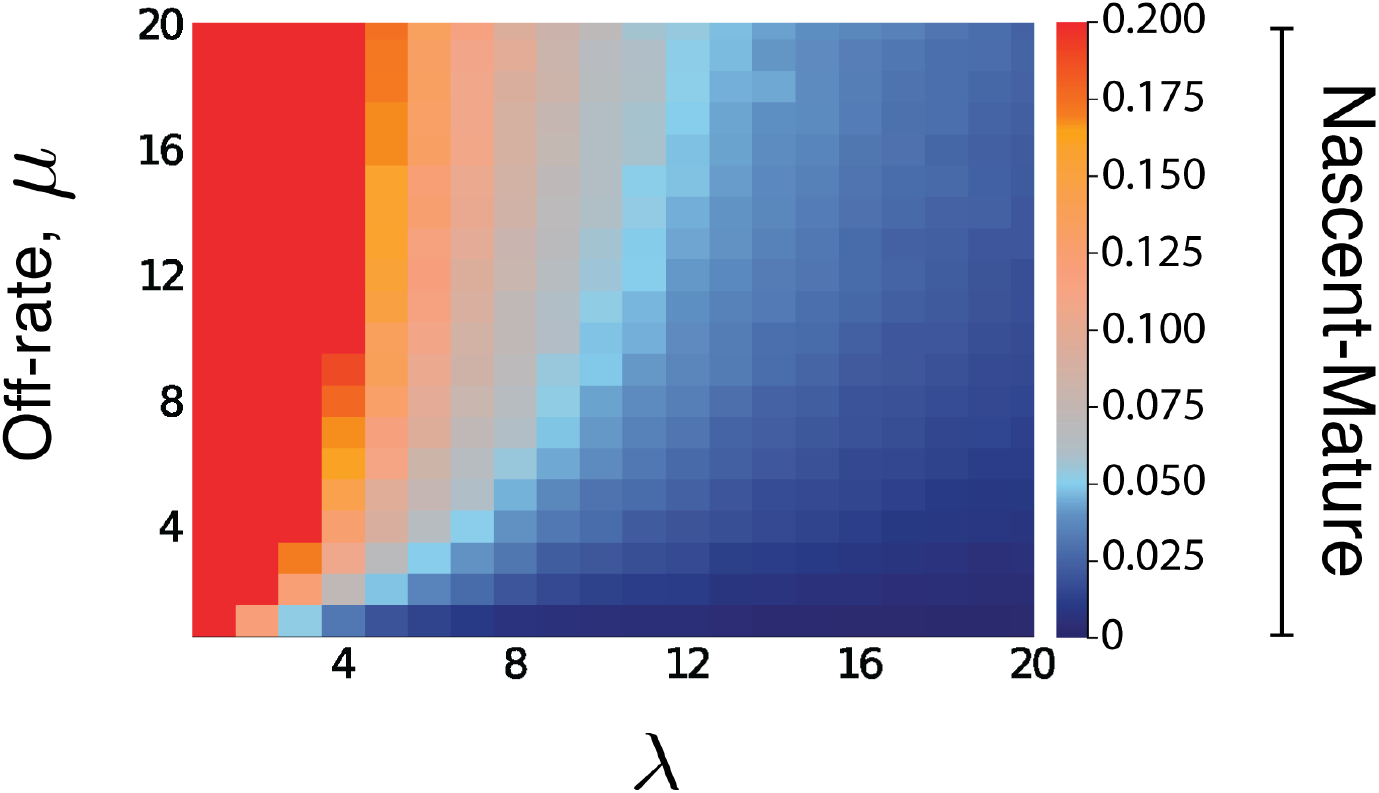
Heatmap for the intrinsic contribution to the covariance for nascent-mature pathway reporters. The nascent-mature reporter concerns only mRNA and so is independent of all protein-related parameters. The heatmap shows the intrinsic contribution for values of *λ* and *µ* between 0.1 and 20, with the same parameter selections for *K*_*N*_, *K*_*M*_ as in Fig. 4 of the main text. Similar simulations for average nascent mRNA levels of 3 and of 8, and mature mRNA levels of 30 and of 160 produced almost identical heatmaps.

Dataset S1

Simulation results of the pathway-reporter method for constitutive genes across 60 different parameter values. We consider noise on all of the parameters except mRNA decay in a constitutive model with mRNA maturation and protein translation. Refer to the excel spreadsheet Constitutive Results.xlsx for full details of the simulation, including the chosen noise distributions and parameters.

Dataset S2

Simulation results for the overshoot estimate in the pathway-reporter method for bursty genes across 448 different parameter values. Refer to the excel spreadsheet NoiseFree Results.xlsx for full details of the simulation, including the chosen noise distributions and parameters.

